# Exploring genomic data coupled with 3D chromatin structures using the WashU Epigenome Browser

**DOI:** 10.1101/2022.01.18.476849

**Authors:** Daofeng Li, Deepak Purushotham, Jessica K. Harrison, Ting Wang

## Abstract

Biological functions are not only encoded by the genome’s sequence but also regulated by its three-dimensional (3D) structure. More and more studies have revealed the importance of 3D chromatin structures in development and diseases; therefore, visualizing the connections between genome sequence, epigenomic dynamics (1D) and the 3D genome becomes a pressing need. The WashU Epigenome Browser introduces a new 3D visualization module to integrate visualization of 1D (such as sequence features, epigenomic data) and 2D data (such as chromosome conformation capture data) with 3D genome structure. Genomic coordinates are encoded in 3D models of the chromosomes; thus, all genomic information displayed on a 1D genome browser can be visualized on a 3D model, supported by genome browser utilities and facilitating interpretation of genomic data. Biological information that is difficult to illustrate in 1D becomes more intuitive when displayed in 3D, providing novel and powerful tools for investigators to hypothesize and understand the connections between biological functions and 3D genome structures.

## Main

Three-dimensional (3D) genomic structures are vital for the regulation of gene expression and cell function^1^. Chromatin conformation capture technologies based on DNA proximity ligation, such as high-throughput chromosome conformation capture (Hi-C)^2^ and chromatin interaction analysis by paired-end tag sequencing (ChIA-PET)^3^, are used to study 3D structures in the genome. These assays usually generate many chromatin interactions as pairs of genomic loci linked by proximity ligations. Based on these chromatin interactions, many algorithms, such as Dip-C/hickit^4^, 3DMax^5^, and LorDG^6^, have been developed to model the 3D genome structure. Large consortia, such as ENCODE^7–9^, Roadmap Epigenomics^10, 11^, IHEC^12^, and 4DN^13^ projects have generated tens of thousands of genome-wide data of transcription factor binding sites and epigenetic marks, including DNA and histone modifications, and chromatin interactions, across cell types and tissues. Biologists wish to visually explore the connections between these vast genome-wide resources and 3D genome structures, which will facilitate generation and testing of diverse hypotheses. This presents a daunting challenge to conventional genome browsers^14, 15^ where most of the genomic data visualization is conducted in the context of linear 1D genomic coordinates.

Interactive visualization of macromolecular structures is beneficial for biological research. Several tools including NGL viewer^16^ and LiteMol^17^ have been developed alongside with special data formats such as PDB and mmCIF file format^18^. A genomic scientist wishing to visualize structural details of the genome would need to convert chromosomal structural data to one of the established formats used by the macromolecular field for visualization. This approach poses a significant limitation - the inability to handle multi-resolution or region-based queries. Genomics 3D structure data modeled with multiple resolutions is difficult to retrofit into established formats. Scientists may also want to look at a specific local region, however the current viewers do not offer functionality to restrict data sources to the requested region. For example, the GSDB^19^ database stores individual structure data in PDB format separately by chromosomes and by resolutions.

The WashU Epigenome Browser was invented in 2011 to bring an interactive tool for exploration of genomic data in a web browser^15^. In this study, we significantly expanded the browser functions to allow investigators to visually explore 1D, 2D and 3D data in a single webpage. The key innovation is to thread the linear genomic coordinates into a multi-resolution 3D model of the genome, which is encoded by a novel data format called g3d. Any genomic information anchored on the linear coordinates can be mapped to and displayed on the 3D model. All typical genome browser functions from the conventional WashU Epigenome Browser, such as query, label, zoom, pan and animation are found in the 3D browser to facilitate exploration of genomic data in the context of a 3D chromatin model. Novel functions specific to 3D data enable users to intuitively examine biological information that is difficult to display on a typical linear genome browser. These novel 3D browser functions provide powerful tools for investigators to form and test hypotheses about biological functions and 3D genome structure.

## Results

### Integrated data visualization using panels on the same webpage

To facilitate direct comparison between the WashU Epigenome linear browser view and new 3D browser view, we implemented a panel system design for integrated data visualization. The linear browser panel is used to display 1D and 2D data, such as genomic sequence, gene annotation, ChIP-seq signal, and Hi-C heatmap, while 3D models are displayed in multiple 3D browser panels (Figure 1). The panels are called ‘responsive panels’, such that each panel can be resized and repositioned using drag-and-drop and maximized to full screen. When resizing or dragging, each panel’s view will automatically resize based on the panel width. The panel layout is saved into sessions, which is a convenient mechanism for sharing between collaborators. By default, regions displayed on the linear browser panel are highlighted on the 3D model in the 3D browser panel. The 3D highlighted region is synched with the linear browser and automatedly updates when users pan, zoom, and reposition on the linear browser (Figure 1B, 1D). The sync function is reciprocal – clicking anywhere on the 3D model will bring up a popup menu. Using this popup menu, users can jump to the selected genomic segment on the linear browser or label the current segment as a particular shape or arrow (Figure 1C). The 3D browser contains two viewing panels: a main viewer and a thumbnail viewer (Figure 1E). The thumbnail viewer is synched with the main viewer and supports a global display when a user focuses on specific local details of the 3D model through the many configurations in the main viewer.

**Figure 1:**
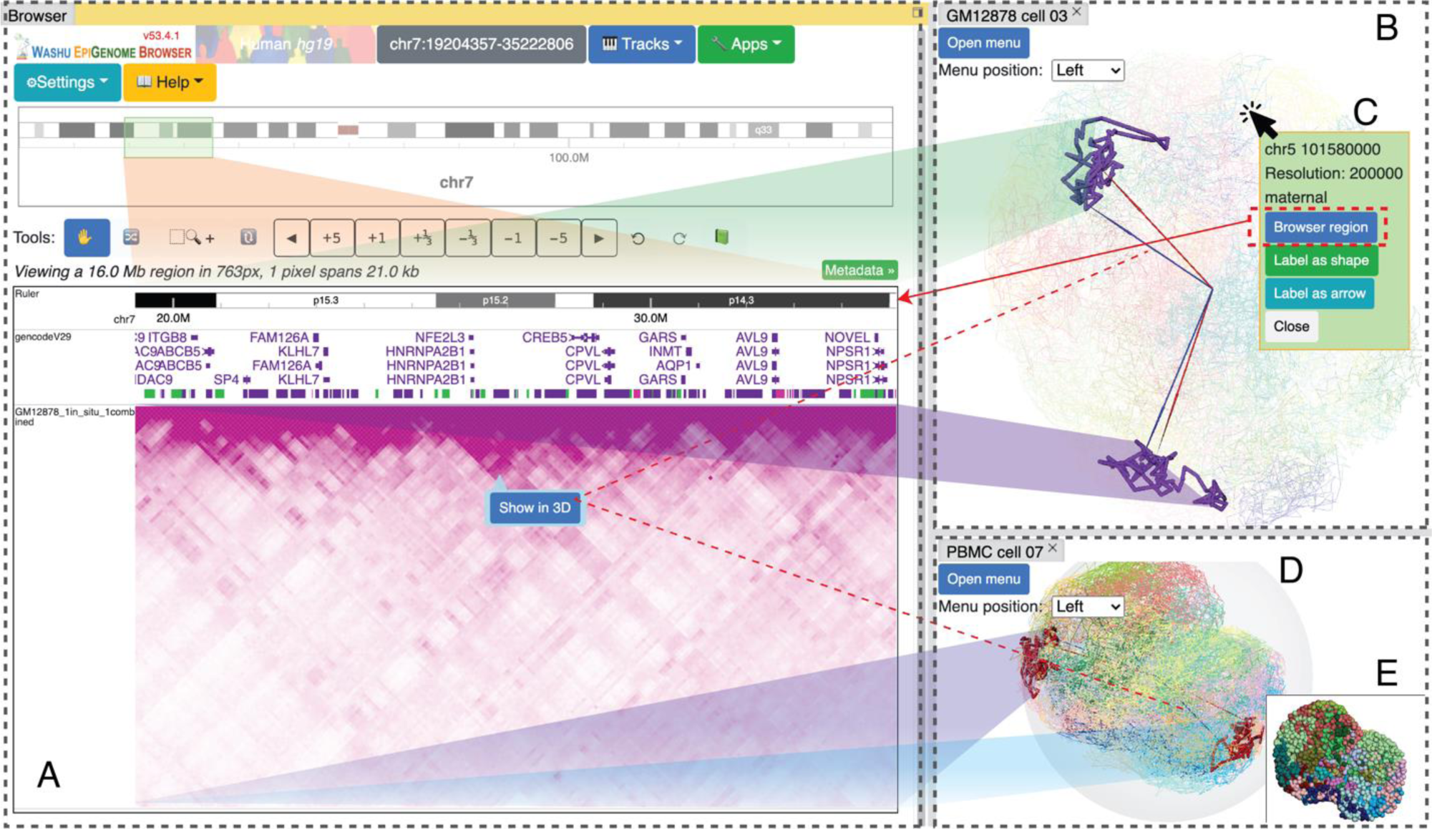
3D visualization integration with panels. The linear genome browser is displayed on the left (A); two 3D structures are displayed on the right (B, D). From the 3D structure, click a 3D segment to reveal a popup menu with options to highlight the linear browser region or label this segment (C). 3D viewer also comes with a thumbnail by default, which view is synchronized with the main 3D viewer (E).

### 3D model – data file, turn and rotate, zoom and pan

3D modeling software/tools usually generate 3D models of chromatin in three-dimensional coordinates (x, y, and z) with 1D chromosomal locations marked along the model. We developed the g3dtools package to convert this 3D model data into g3d format, which supports remote range queries, multiple resolutions, haplotypes and cell/sample types. The g3d format allows data to be compressed internally for efficient storage and fast transfer. The WashU Epigenome Browser uses a JavaScript package, g3djs (https://www.npmjs.com/package/g3djs), to access g3d files and load the 3D model. Each genomic region with 3D coordinates is treated as an ‘atom’, and nearby atoms/regions are connected (as edges) when the 3D model is viewed in cartoon or line style. Once the 3D model is loaded in the browser, users can use the mouse or touchpad to turn, rotate, zoom and pan the model in a manner identical to how protein or other macromolecular structures are typically visualized. A complete list of supported mouse controls as well as a full tutorial are included in Supplemental Notes. A series of video tutorials illustrating 3D browser operations can be found at https://bit.ly/eg3dtutorial.

### 3D visualization configurations

Users are provided with many utilities to control and configure the display of data in 3D by using an interactive menu. The menu is divided into seven sections, including configuration of model data, viewer layout, highlighting and labeling, numerical painting, annotation painting, animation, and export functions (‘3D viewer menu’ section in Supplemental Notes). The model data section contains the resolution selection and enables toggling between models, either organized by haplotypes or individual cells. The layout section contains options for the 3D main and thumbnail viewer layout and the thumbnail structure display style. The highlighting & labeling section supports region highlighting by genomic regions or genes. Users can submit a gene symbol or genomic region for labeling on the 3D model or upload a text file that contains a list of regions/gene symbols for batch labeling. Users can customize the label style including color, text, and shape; labels can also be shown as arrows pointing to the desired region/gene and be customized for color and style (Figure 2, video tutorial).

**Figure 2:**
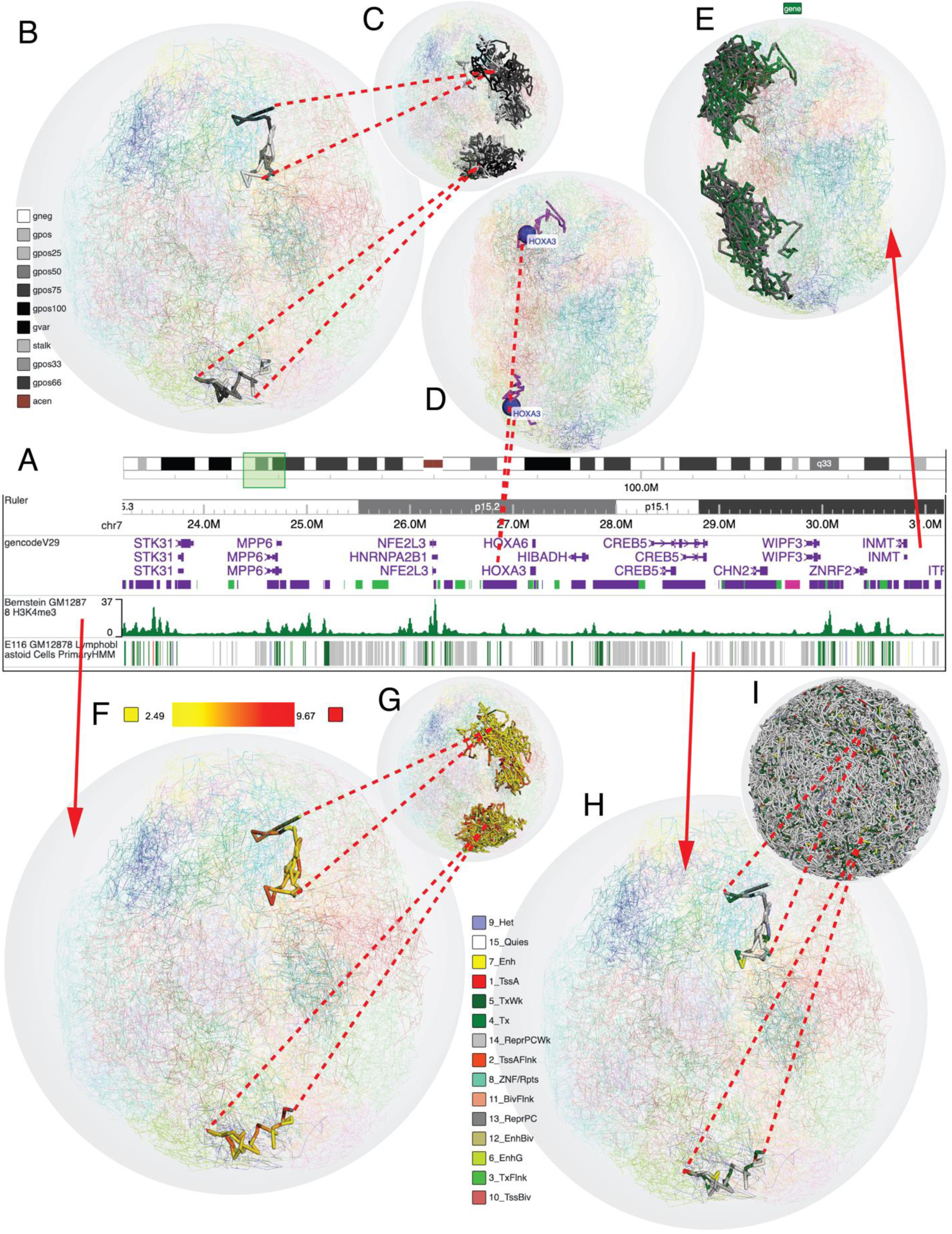
Highlighting and decoration in 3D structure. Browser view is showing cytobands, gene annotations, H3K4me3 signal and chromHMM segmental annotation for GM12878 (A). Cytoband painting in 3D of current browser region (B) and current chromosome which is chr7 (C), please note the 3D model contains both models from the two haplotypes (paternal and maternal); thus, 2 regions and chromosomes are displayed (same applied to following contents). One gene, *HOXA3*, is highlighted in the 3D model (D). Gene annotation is used to paint the 3D structure; green indicates gene location (E), clicking the ‘gene’ button can customize the highlighting color. H3K4me3 signal is used to paint the 3D model in current region (F) and chromosome (G). Highlighting colors and scale can be customized from the legend and configuration menu. ChromHMM annotation is used to paint the 3D model in current region (H) and chromosome (I).

In addition to labeling specific regions and genes on the 3D model, the 3D browser allows users to decorate or paint the 3D model using genomic data. Painting is achieved by drawing a tube over the lines that represent the underlying chromatin. There are two general painting styles: annotation and numerical. In annotation painting, users can decorate the 3D model using any genomic annotation in a segmentation type format, such as cytoband (Figure 2B, 2C), gene annotation (Figure 2D, 2E), and chromHMM segmentation (Figure 2H, 2I). The same function makes it easy to paint compartmental calls, such as topological associated domain calls, which have a customizable format. Genes from the linear browser track can be directly painted on the 3D chromatin (Figure 2D), including painting all genes (Figure 2E). Users can also customize colors of any annotation category to achieve the desired visualization effect (tutorial in Supplemental Notes and videos).

In numerical painting, users can decorate the 3D model with numerical data, such as GC content or ChIP-seq signals, in color gradients. Numerical painting decorates the 3D structures using any browser loaded or user-provided bigwig file, or gene expression table with RPKM/FPKM values. Users can configure paint thickness and background line opacity. Users can also configure the numerical scale and colors to adjust the color gradient (Figure 2F, 2G, and video tutorial).

Both annotation and numerical painting can be done simultaneously at region, chromosome and genome-wide scale. In addition, the animation function can iterate multiple structures or models and display them as an animation. This function can be useful for visualizing dynamic changes of 3D data across a time series (video tutorial). The export function can export the 3D structure as a high-quality figure for presentation purposes.

### Visualizing compartments, chromatin loops and TADs

The 3D browser allows for a straightforward view of the two major spatial compartments, A and B compartments, which correspond to active and inactive chromatin, respectively (Figure 3). Multiple compartment call formats are supported. Typically, researchers use positive values for A and negative values for B compartment, and the data is usually displayed as a wiggle track in a linear genome browser (Figure 3B, F, G). Figure 3A displays the genome-wide compartment calls in GM12878 cell^20^ at 200KB resolution. The B compartments are physically close to the nuclear membrane, whereas the A compartments are inside the nuclei – the physical separation between A/B compartments can be much more easily appreciated in a 3D model. Zooming-in reveals detailed compartmentation at a chromosome (Figure 3C) and region level (Figure 3H, I) at 20KB resolution. For example, the *HOXA* cluster is within the active chromatin compartment as seen from the wiggle track (Figure 3G) and 3D view (Figure 3I). Another compartment definition contains 5 subtypes (A1, A2, B1, B2 and B3); this format can also be viewed as an annotation track in a linear browser (Figure 4A) and in a 3D view (Figure 4E). The compartmental data on the 3D browser can be customized according to compartment scores and category using color and gradient/scales (Figure 3D, E; 4E). The 3D browser also makes viewing chromatin loop and topologically associated domains (TADs) intuitive (Figure 4). Users can view Hi-C loop anchors along with the loops in 3D (Figure 4A-D). Genomic segments inside of the loop are labeled in a different color corresponding to the ‘Domain’ track. Compartmental annotation can be used to paint the 3D structure (Figure 4E), and the color of each compartment can be customized (Figure 4G). Users can also load genomic locations of TADs to paint the 3D structure (Figure 4H-L).

**Figure 3:**
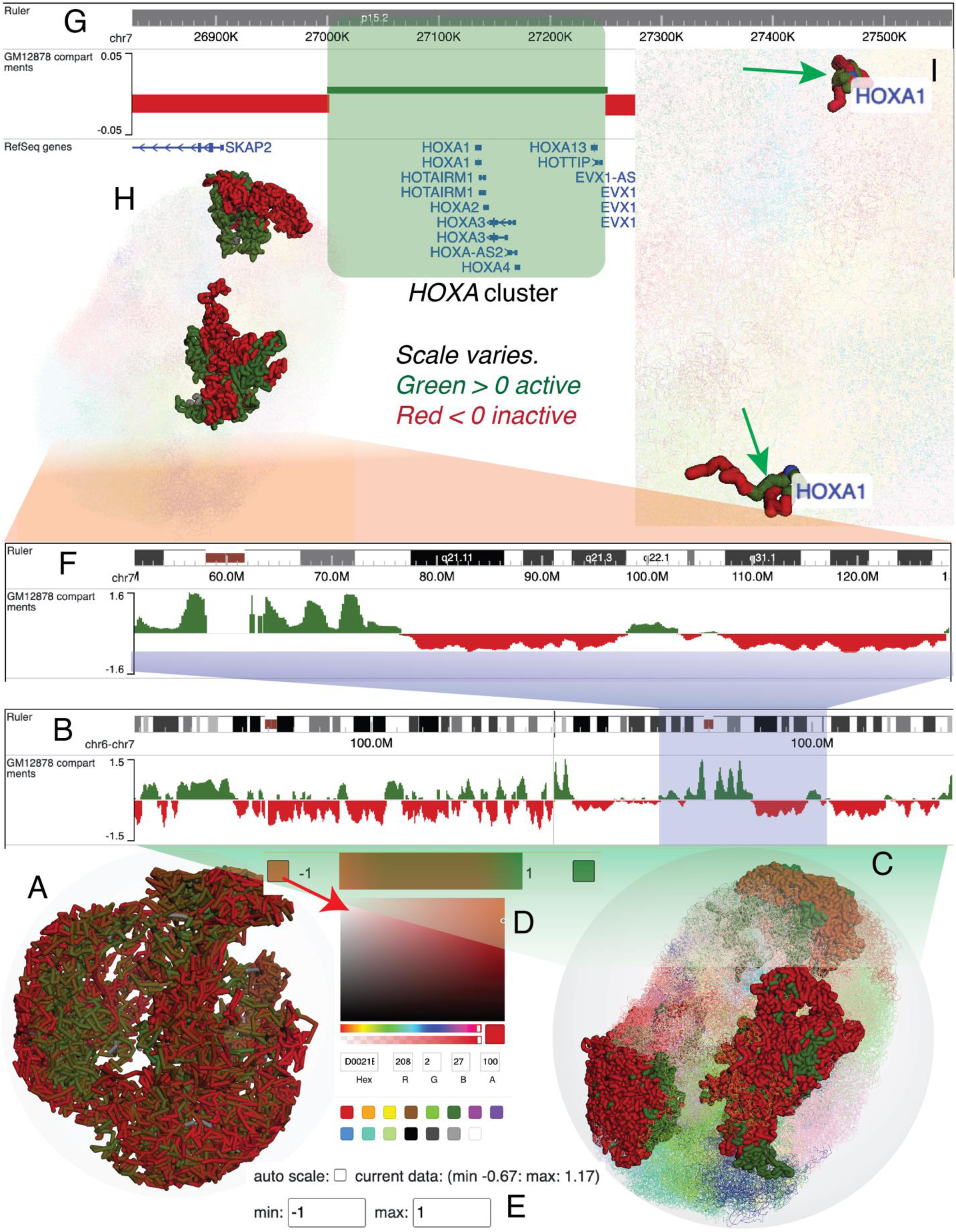
Compartment visualization in 3D structure. Genome wide compartment annotation of GM12878, where green indicates action chromatin and red indicates inactive/silent chromatin at 200KB resolution, where we can observe green (active) chromatin tends to exist inside nuclear while red (inactive) chromatin exists closer to the nuclear envelop (A). Zooming into whole chromosome 6 and 7 showing the compartment score in browser as wiggle track (B), positive values (green) and negative values (red) indicate active and inactive chromatin, respectively. 3D model of current chromosome 6 and 7 (C) at 20KB resolution. Color for the gradient and scale can be changed from legend and configure menu (D, E). Continue zooming to a region in chr7 showing the compartment signal in browser (F) and in 3D (H). Zooming into the region around *HOXA* cluster, where we can tell *HOXA* cluster is in the active chromatin (G, I), of which *HOXA1* gene is displayed as a blue sphere around with green (active) chromatin (I).

**Figure 4:**
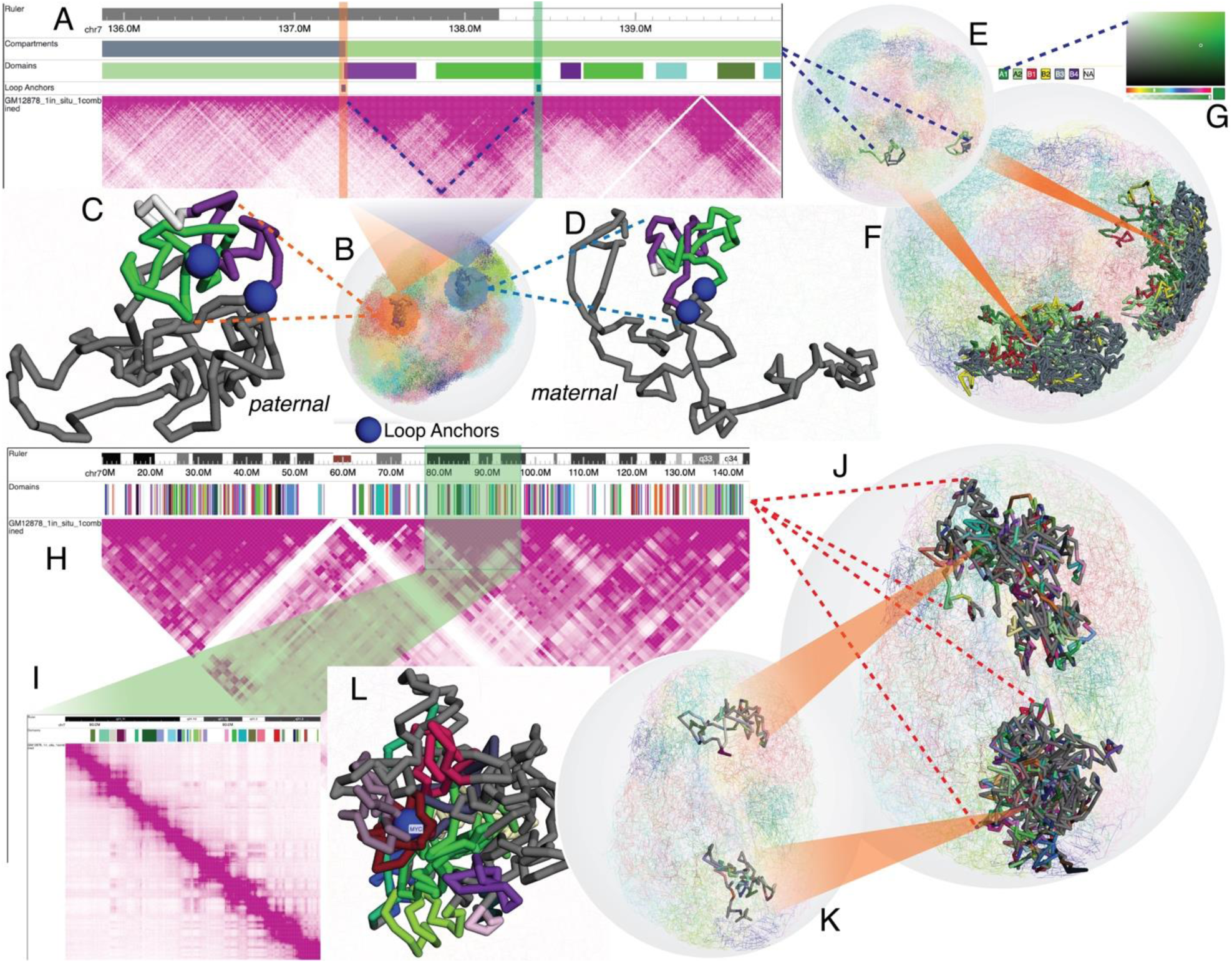
Viewing chromatin loops and TADs. Typical genome browser view with ruler, compartments annotation with subtypes, bed track with loop anchor locations and Hi-C track for GM12878 (A). Blue dashed line indicates the domain/loop, which is highlighted in B, C and D. The orange and green rectangles indicate loop anchors. 2 sub-domain/loops colored purple and green in the browser view are shown as well in the 3D view with the same color. Blue spheres in the 3D model indicate same loop anchors as in the browser view. Compartmental annotation is used to paint the 3D model in current region (E) and chromosome (F). Color of each subtype can be customized after clicking each type of button in the legend (G). Browser view with a ruler, domain annotation as bedcolor track and Hi-C track (H). Zoomed view of one domain with TADs annotation and Hi-C map (I). Current domain annotation is used to paint the 3D model in current region (K) and chromosome (J). An example of nearby domain painting of the *MYC* gene (L).

### Use cases

To illustrate the power of using the 3D browser to investigate biological hypotheses, we used the 3D browser to recapitulate results from several published studies.

It was reported that structural differences exist between maternal and paternal alleles^4^. Here we show that the 3D chromatin structures of the two X chromosomes from paternal and maternal alleles are quite distinct and agree with the corresponding haplotype resolved contact maps (Fig 5A-C). The maternal chromosome appears to be more relaxed and the super-loop anchors are more distantly positioned, whereas the paternal chromosome is much denser and the super-loop anchors are adjacent to one another. Many of the loops (indicated by green circle) from the parental Hi-C map are missing from the maternal map, which may translate into the 3D structure differences (Figure 5A).

**Figure 5:**
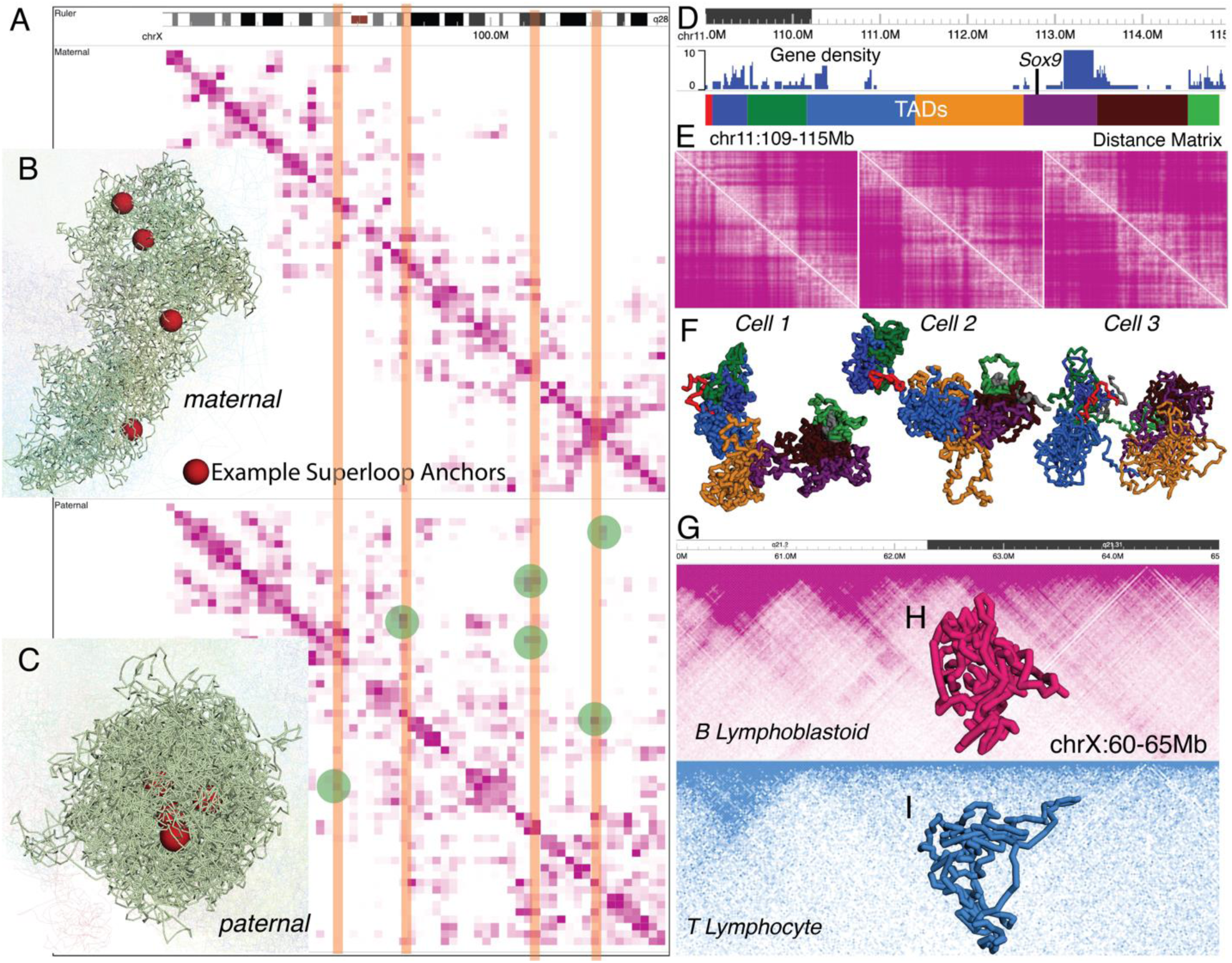
Use cases. The 3D browser is used to recapitulate serval findings from published studies. Haplotype-resolved Hi-C contact maps are shown in the linear browser (A), maternal (top) and paternal (bottom). 3D models from both alleles are displayed (maternal, B; paternal, C). Red spheres indicate example super-loop anchors, which are also highlighted in orange box in the browser view. Green circles represent the loops only presented in paternal allele but not maternal allene. Browser view around mouse *Sox9* locus showing gene track in density mode and TADs annotation (D). Distance matrix and three cells (E) and corresponding 3D models (F). 3D models are colored with TADs annotation. Hi-C contact maps are shown in browser for B lymphoblastoid (blue) and T lymphocyte (pink) (G) at chrX 60-65Mb region, and the corresponding 3D structures (H, I).

Single-cell data has revealed that the 3D genome conformations of individual cells display high levels of variability^21^. We can observe such variability in the single-cell contact maps^22^. This observation can be recapitulated by the 3D models of single-molecule conformations (Figure 5F) with their corresponding single-cell contact maps (Figure 5E) of three individual mouse cells. These distinct 3D models display a 6 Mb genomic region containing the *Sox9* gene and are labeled with TADs^23^ (Figure 5D).

Finally, chromatin structures can be cell type-specific, reflecting a tight association with cell type-specific environment or gene regulation^4^. A broadly defined epigenetic difference between B lymphoblastoid and T lymphocyte can be visualized in 1D (compartmentalization, Figure 5G), 2D (Hi-C contact map, Figure 5G), and 3D models (Figure 5H, I) in parallel, as we illustrate genomic regions 60-65Mb of chrX^4^.

### Data hub for 3D structures

We collected 11,045 3D models from published studies^4, 24–28^, converted them to g3d format and built data hubs. Most of the 3D models were generated from single-cell studies from species including mouse, human, yeast and *P. falciparum* (See Supplemental Table 1 for details). The data hubs can be loaded from each corresponding genome assembly main page.

## Discussion

WashU Epigenome Browser is among the first tools^29^ to display 1D, 2D and 3D genomic data in a single webpage (video tutorials in YouTube: https://bit.ly/eg3dtutorial). The fundamental innovation is how we conceptualize the usage of genomic coordinates. All classic genome browsers anchor genomic data on a one-dimensional, straight-line axis, emulating a process of untangling, straightening, and stretching genomic DNA. This process obviously makes it both convenient and straightforward to overlay genomic data on this linear DNA, nevertheless, it destroys the spatial configuration of the chromatin, or the 3D structure of the underlying genomic DNA. What we strive to achieve is to maintain the spatial configuration by threading the linear genomic coordinates through a 3D structure model. In another word, we simply provide a new coordinate system that combines the linear genome location and the 3D spatial location and build visualization tools on top of this new coordinate system. Recent advancements in genomic technologies and computational algorithms have provided unprecedented opportunities to probe chromatin interactions and generate 3D models of chromosomes at high resolution. These models not only facilitate the investigation into the formation and function of the 3D genome, but also provide a new paradigm to display and interact with genomic data. The new WashU Epigenome Browser 3D viewer enables investigators to intuitively examine 3D features, such as loops and TADs, to visualize all genomic data on 3D genome coordinates, and to explore the dynamics of the 3D genome structure. It will become a valuable tool in genomics.

## Methods

### The g3d file format and command line tool: g3dtools

The g3d file format uses a header and body design. The header section is a fixed 64Kb block, and the body utilizes MessagePack (https://msgpack.org) for efficient binary serialization and Zlib (https://zlib.net/) for data compression. Structural data for each resolution/category is saved in one binary block. We provide Python and JavaScript API for querying a g3d format file. Users can easily query data from a g3d file based on a specific region, chromosome, or the entire genome. A command-line tool called ‘g3dtools’ is provided to generate a file in g3d format. The tool g3dtools can be easily installed using the Python Package Installer. Users can customize aggregation settings (aggregate the data to lower resolution by applying scales) in g3dtools to convert 3D structure in text format to the binary g3d format. For example, when converting data at 20Kb resolution to 40Kb resolution, g3dtools will use the center point of 2 nearby 20Kb regions, likewise for 60Kb, 80Kb resolutions, etc. Users can also provide a 3D structure model based on different resolutions. G3dtools will then index and combine them into one single g3d file. G3dtools also supports other 3D genomic structure formats such as nucle3d or formats generated by other tools such as pastis and Dip-C/hickit.

### g3d file visualization

3D genomic structure visualization based on the g3d file was developed using 3Dmol.js package. We wrote a customized parser for supporting g3d format in 3Dmol.js, where each chromosome bin is treated as an atom and only nearby atoms are connected/bonded.

### Code availability

g3d related source code is freely available at https://github.com/lidaof/g3d, and g3dtools documentation is available at https://g3d.readthedocs.io/en/latest/. The Browser codebase is available at https://github.com/lidaof/eg-react, and documentation can be found at https://eg.readthedocs.io/en/latest/. This repository https://github.com/lidaof/eg-3d-demo contains demo files for 3D visualization and brief instructions. All code is open source.

### Data availability

All data used in this study is published or public from open databases, they are listed in the table below:

| Data set | Reference or URL |
| --- | --- |
| GM12878, T lymphocyte HiC | <sup>20</sup> |
| GM12878, PBMC 3D models | <sup>4</sup> |
| GM12878 H3K4me3 | <sup>8</sup> |
| GM12878 chromHMM | 11 |
| GM12878 compartments signals | https://data.4dnucleome.org/files-processed/4DNFILYQ1PAY/ |
| GM12878 subtypes compartments | 20 |
| GM12878 loops, TADs annotation | 20 |
| Mouse 3D model, distance matrix | 22 |
| Mouse TADs annotation | 23 |

## Acknowledgements

We thank the 3Dmol.js team (https://3dmol.csb.pitt.edu/) for the fantastic 3D graphics library. We thank David Sehnal for suggestions on data structure of g3d format, Longzhi Tan and Luca Fiorillo for the data discussion. We also thank all the members in Wang laboratory for testing and helpful feedbacks. This work was supported by NIH R01HG007175, U01CA200060, U24ES026699, U01HG009391, UM1HG011585, U41HG010972, U24HG012070.

## Author Contributions

T.W. conceived and oversaw all aspects of the study, supervised research and provided constructive feedback. D.L. developed and is maintaining the 3D visualization module in the Browser. D.P. and J.K.H. provided key contributions and useful feedback. The manuscript was written by D.L. and T.W. with additional contributions and edits from D.P. and J.K.H.

