## Supplemental notes for "Exploring genomic data coupled with 3D chromatin structures using the WashU Epigenome Browser"

### Collected 3D data hubs

**Supplemental Table 1: 3D data statistics collected for each genome assembly.**

| Genome | Count | GEO accession or project link | Reference |
|---|---|---|---|
| Human (*hg19*) | 34 | GSE117109 | [1] |
| Mouse (*mm10*) | 10 | GSE117109 | [1] |
| Mouse (*mm10*) | 1227 | GSE121791 | [2] |
| Mouse (*mm10*) | 9770 | GSE162511 | [3] |
| Yeast (*sacCer3*) | 1 | https://noble.gs.washington.edu/proj/yeast-architecture | [4] |
| P. falciparum (*pfal3d7*) | 3 | https://noble.gs.washington.edu/proj/plasmo3d/ | [5] |

### Supported mouse controls on the 3D model

The 3D browser utilized 3dmol.js package to display the 3D model based on a custom data parser; thus, the mouse controls of 3dmol are also supported and are listed below (from 3dmol.js documentation):

| Movement | Mouse Input | Touch Input |
|---|---|---|
| Rotation | Primary Mouse Button | Single touch |
| Translation | Middle Mouse Button or Ctrl+Primary | Triple touch |
| Zoom | Scroll Wheel or Second Mouse Button or Shift+Primary | Pinch (double touch) |
| Slab | Ctrl+Second | Not Available |

### The g3d file format

`g3d` is a new format we developed for visualizing 3D structural data on the WashU Epigenome Browser. `g3d` format is a binary format based on the bed-like format of data and contains x, y, z coordinates for each genomic bin. Documentations for how to prepare a `g3d` file is available at g3dtools documentation.

`g3dtools` is a Python package we developed to generate the binary `g3d` files. Such as the example input below, which is a 6 column bed-like text file (6th column is optional):

```
chr7    16760000    -14.3866688728  -36.3919302029  19.8483965881   m
chr7    16760000    -24.9116268071  50.0521268287   9.91073185128   p
chr7    25160000    -10.1170055526  -34.8975763469  20.2401719179   m
chr7    25160000    -21.8210915649  27.1128556621   13.4856945965   p
chr7    33540000    -4.11059384846  -54.4940083464  4.21321135564   m
chr7    33540000    -12.0040359857  31.5960497183   26.6925954134   p
chr7    41940000    5.75342635105   -55.4976428728  8.65307697332   m
chr7    41940000    -23.7372022413  36.0614692267   31.919119243    p
chr7    50320000    -10.7099779927  -38.0214001171  25.8308473821   m
chr7    50320000    -28.5142098162  26.6468499001   28.8634805533   p
chr7    26200000    -11.5800097945  -37.9903257744  16.2461100893   m
chr7    26200000    -15.9552426623  27.016940724    17.5722080595   p
chr7    27260000    -14.1883124179  -44.7860807973  12.4104162757   m
```

```
chr7    27260000    -20.0857754297  30.9204143041   18.4774635708   p
chr7    28300000    -18.0160836669  -39.398544495   12.811858164    m
chr7    28300000    -14.9383020843  39.1464516779   17.3743509519   p
chr7    29360000    -11.8032470923  -47.3595095319  13.2828128833   m
chr7    29360000    -12.2445277916  41.2431968179   14.8844908717   p
chr7    30400000    -12.8674349856  -45.0752589744  9.15498568359   m
```

columns are:

- chromosome
- start position
- X (coordinates in 3D)
- Y
- Z
- category (optional)- usually haplotype, cell/sample type or time point information
    - for cell identifier, user can also choose cell-1, cell-2 etc.
    - if omitted, 'shared' will be used
    - example for haplotype:
        - m for maternal
        - p for paternal
        - s for shared

The `g3d` input looks like [3dg format]() except we put haplotype or category info on last column.

The command below can be used to generate a new `g3d` file using the format listed above:

```
g3dtools load test.g3d.bed.gz -o test -s 2,3,4,5,6,7,8,9,10 -n GM12878 -g
hg19
```

#### Tutorial: Interactively using the 3D visualization module on WashU Epigenome Browser

In this tutorial, we introduce how to interactively use the 3D visualization module on WashU Epigenome Browser.

The `g3d` files can be submitted as custom tracks from `Tracks -> Custom Tracks` or by using a datahub. Submitting a `g3d` track will trigger a new panel to open in the browser, which also contains a menu allowing you to customize the visualization, such as change resolution and painting the 3D structure using bigwig (ex. GC percentage) or compartment annotations.

**Submit a g3d track**

From the *Tracks* menu, choose *Remote Tracks*, *g3d* track type and paste the `g3d` file url. You can also input a track label, which is optional.

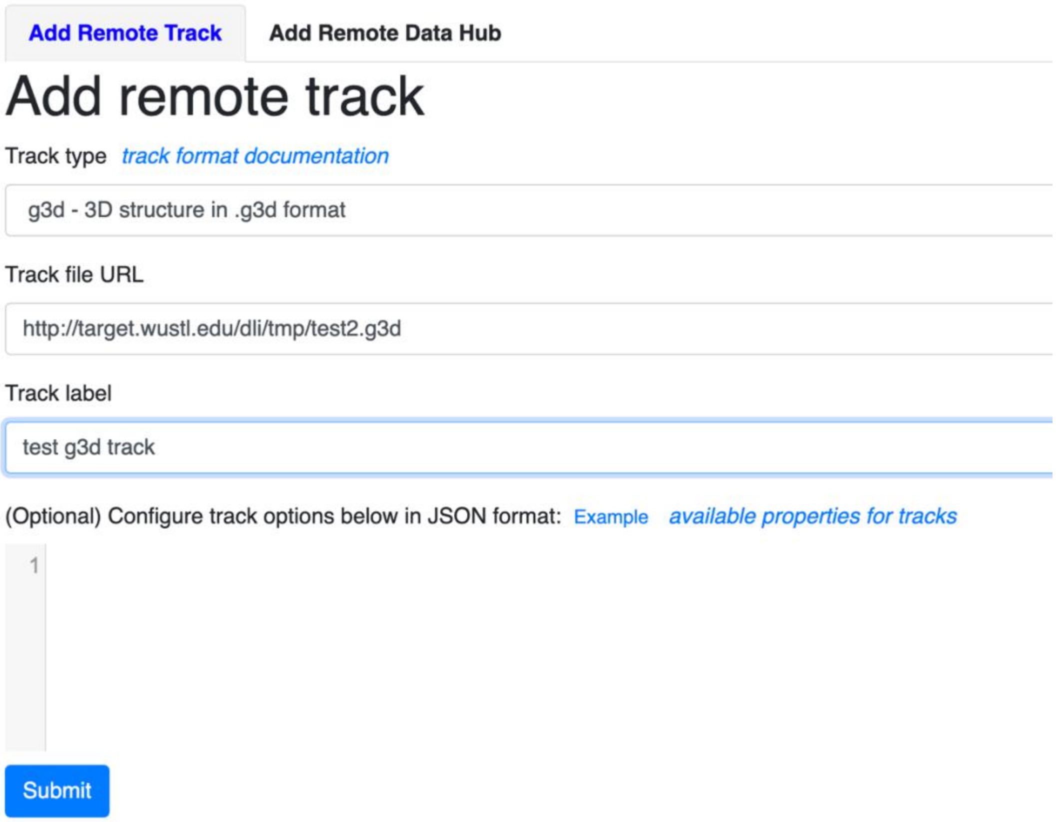

click the *Submit* button, close the Remote track panel, this is how it looks like with default view:

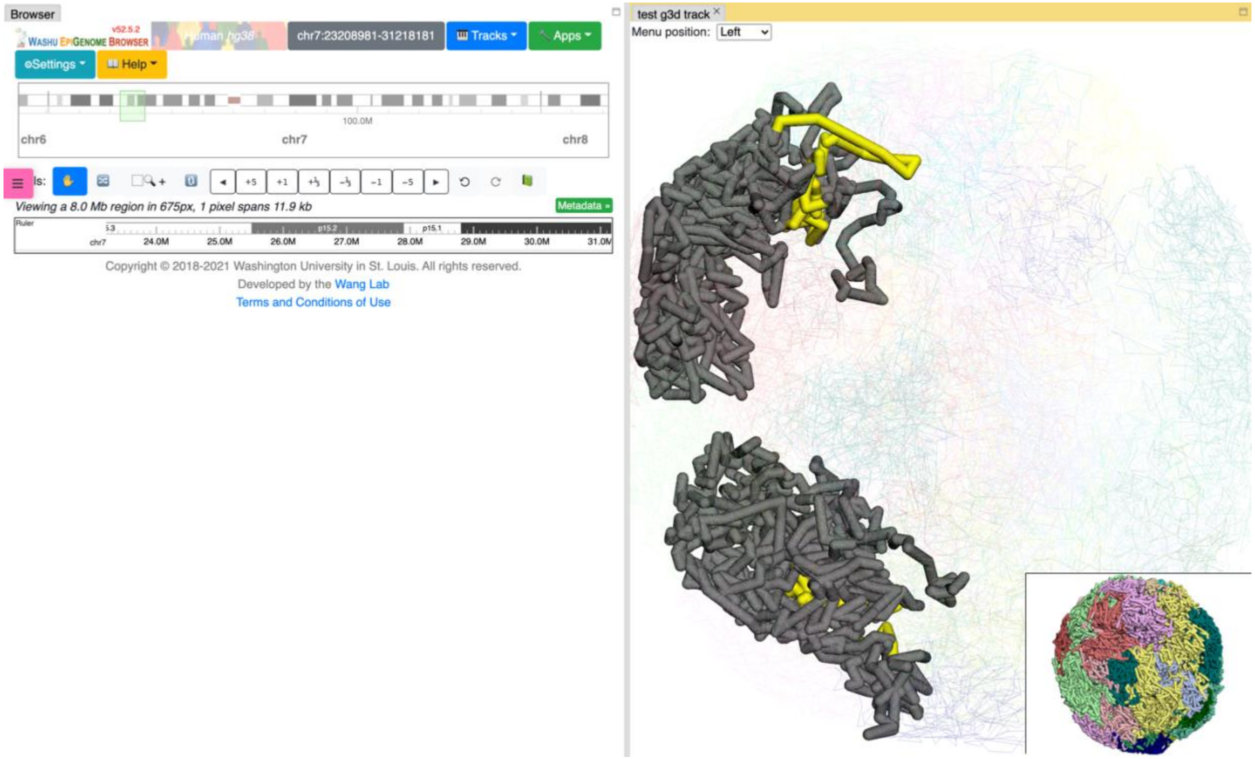

By default, the 3D viewer contains the main and thumbnail viewers, which are synchronized when either of the viewers are zoomed or rotated. The yellow highlighted region indicates the region shown on the linear browser, which is synchronized with the 3D viewer. By changing the linear browser region, the highlighted region in yellow in the 3D viewer will change accordingly.

**3D viewer menu**

Clicking the 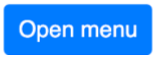

button will open the configuration menu for the 3D viewer, which by default floats to the left of the screen. The menu is grouped to control the model data, layout, highlighting & labeling, painting, animation and export. Each group can be clicked to toggle expansion.

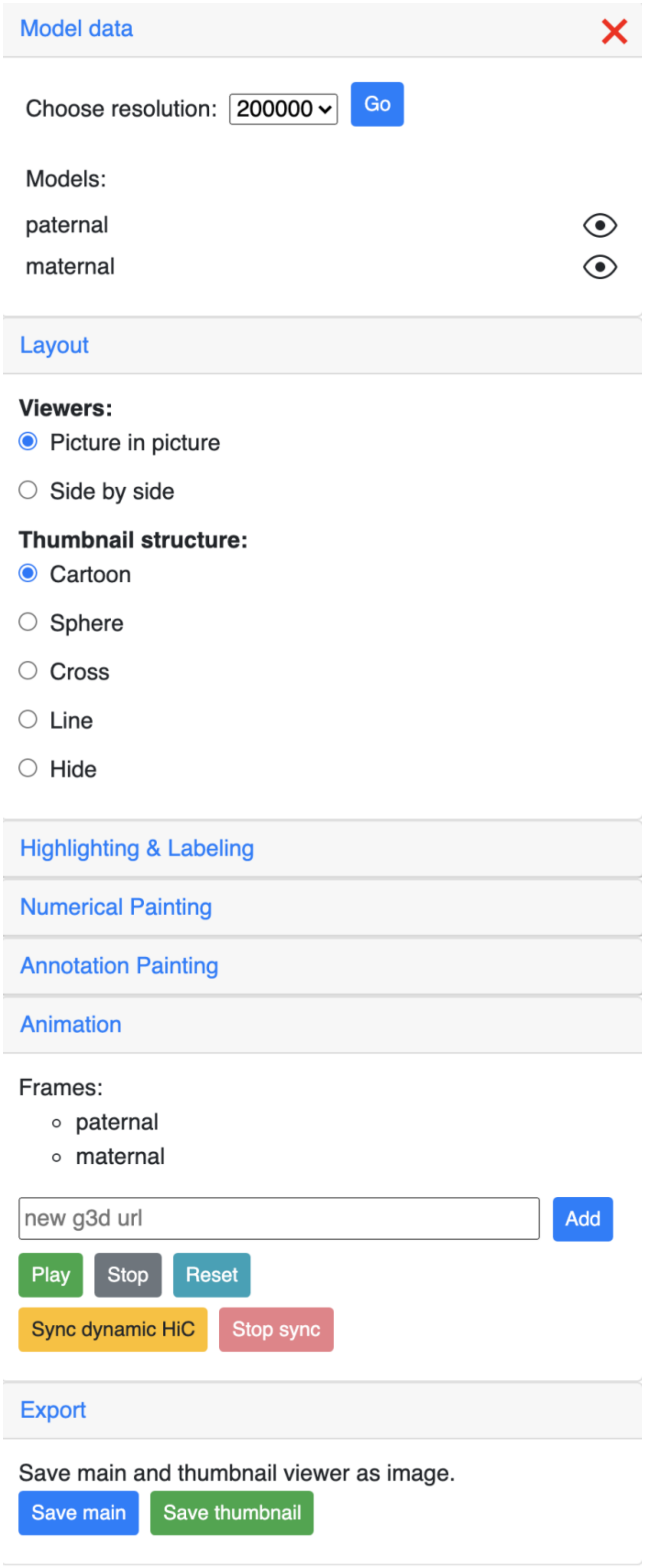
Model data
Choose resolution: 200000
Go
Models:
paternal
maternal
Layout
Viewers:
Picture in picture
Side by side
Thumbnail structure:
Cartoon
Sphere
Cross
Line
Hide
Highlighting & Labeling
Numerical Painting
Annotation Painting
Animation
Frames:
paternal
maternal
new g3d url
Add
Play
Stop
Reset
Sync dynamic HiC
Stop sync
Export
Save main and thumbnail viewer as image.
Save main
Save thumbnail

The menu icon position can also be adjusted using the dropdown menu on the 3D viewer:

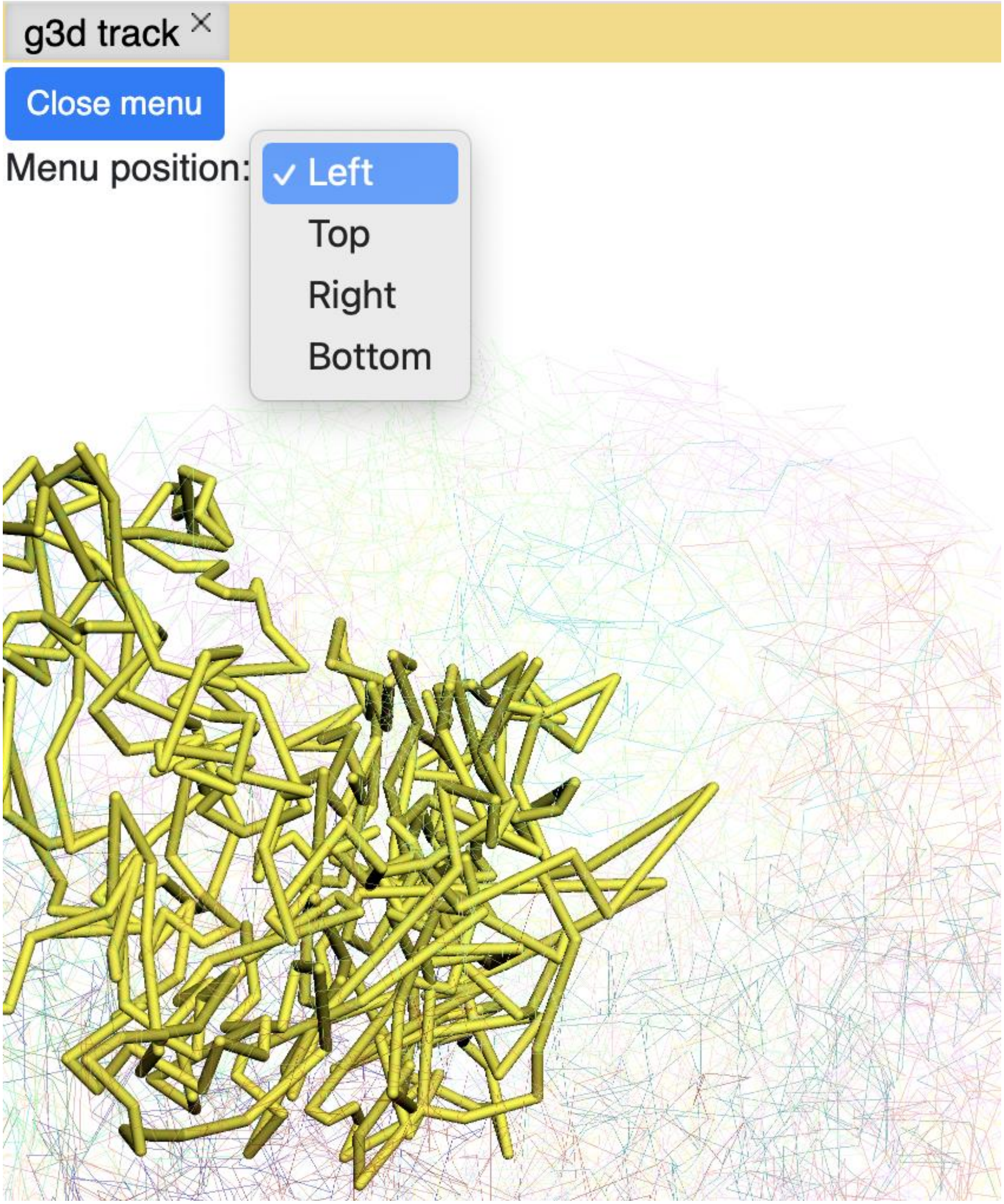

**Config 3D model data**

The *Model data* section can control the resolution of the g3d data.

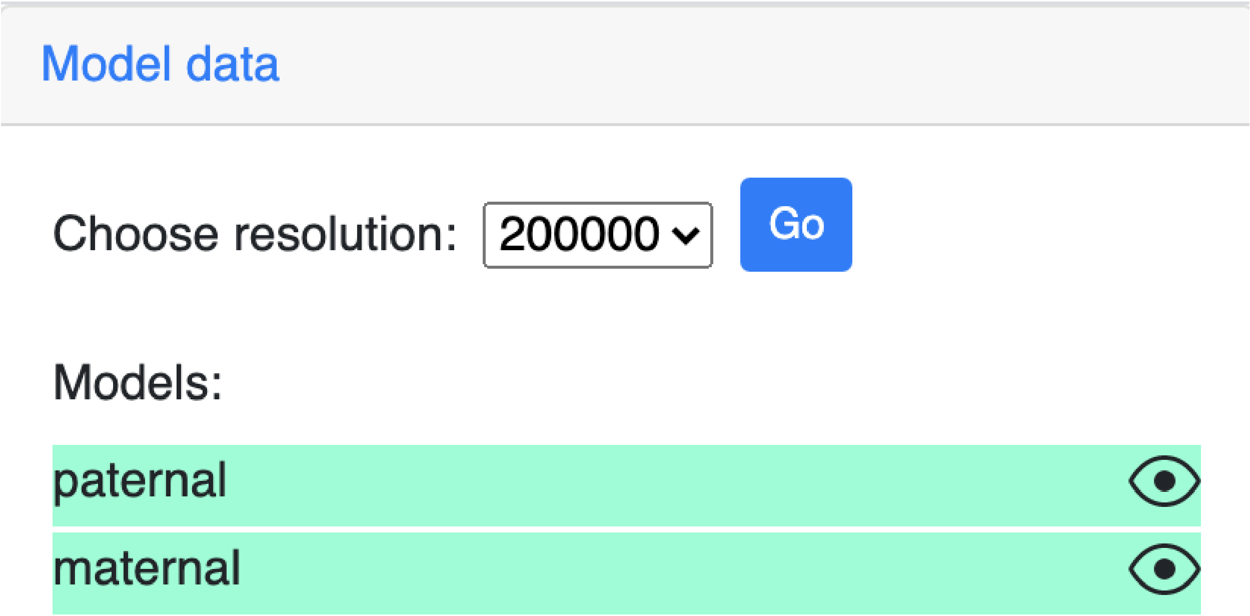

All the models in the g3d file will be listed here and can be displayed or hidden by clicking the eye graphic. For example, the screenshot below indicates the *maternal* model is hidden as seen by the line over the eye graphic:

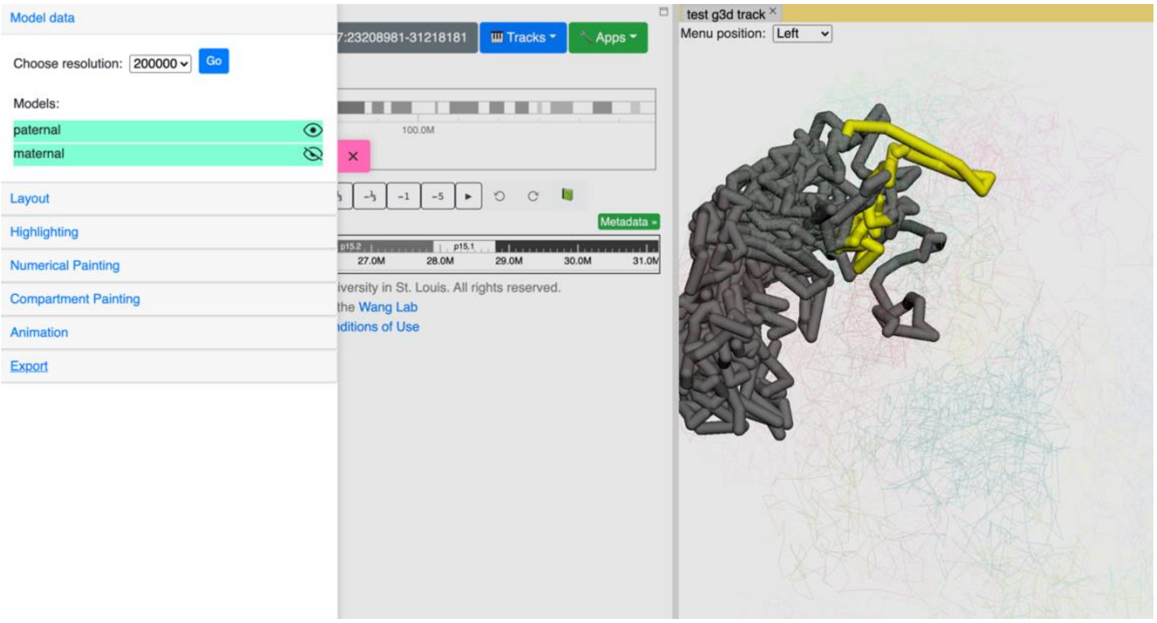

**Config 3D viewer layout**

The *Layout* section is used to control the layout of main and thumbnail viewers.

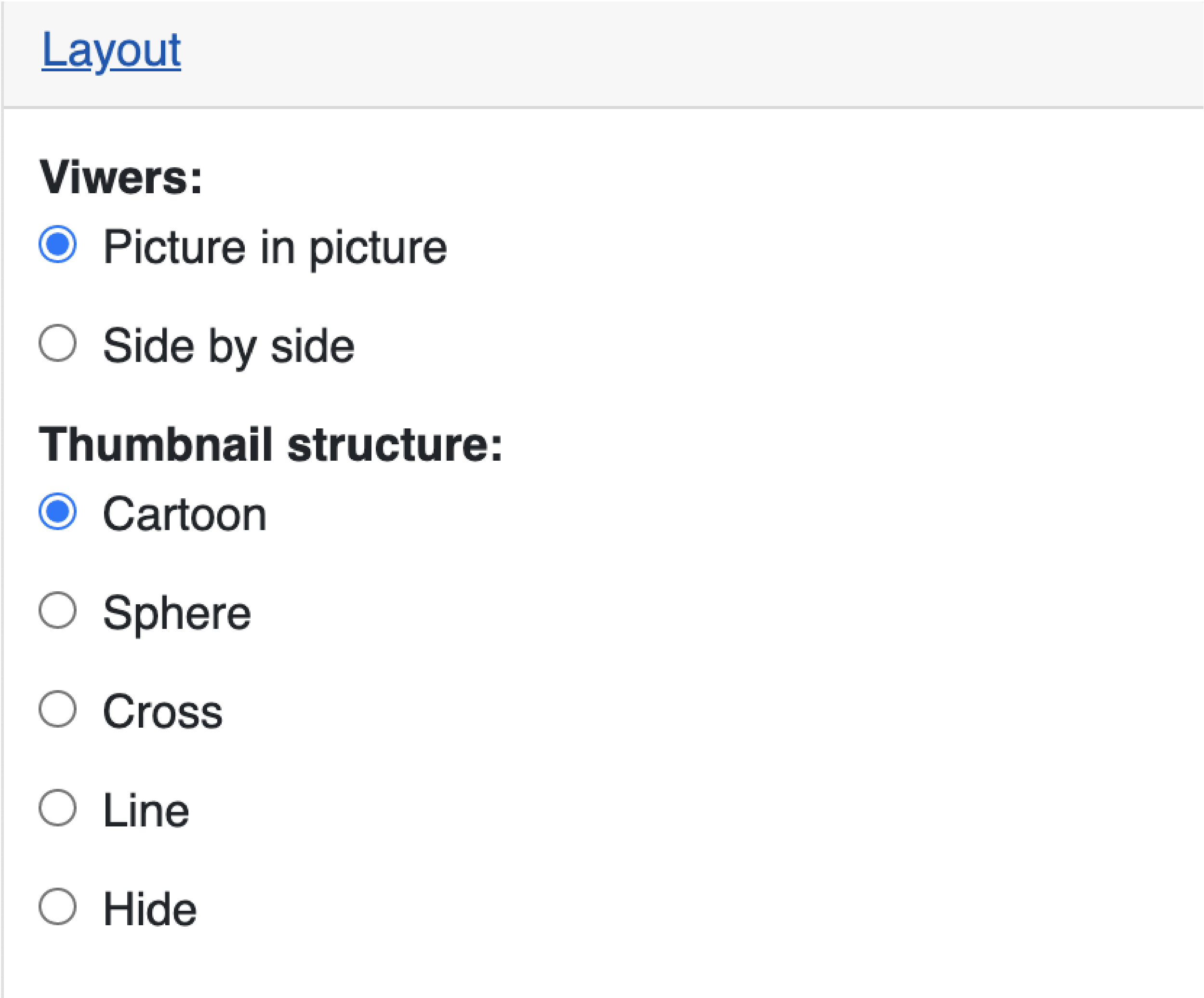

Change the view layout to *side by side*:

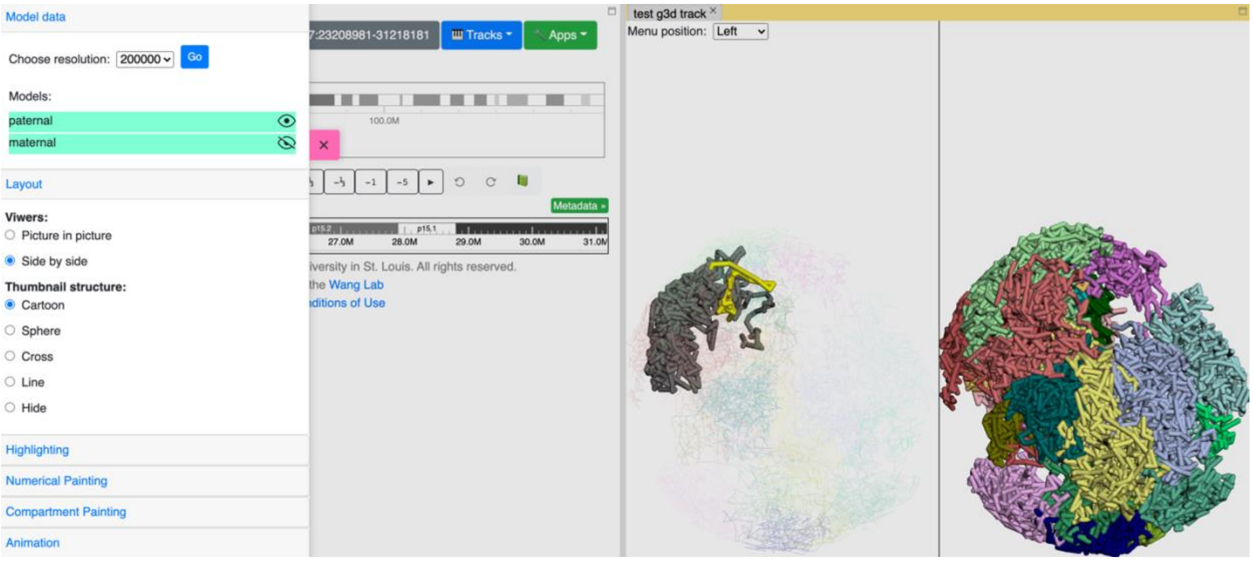

You can also change how the thumbnail structure looks as seen in the *sphere* style below:

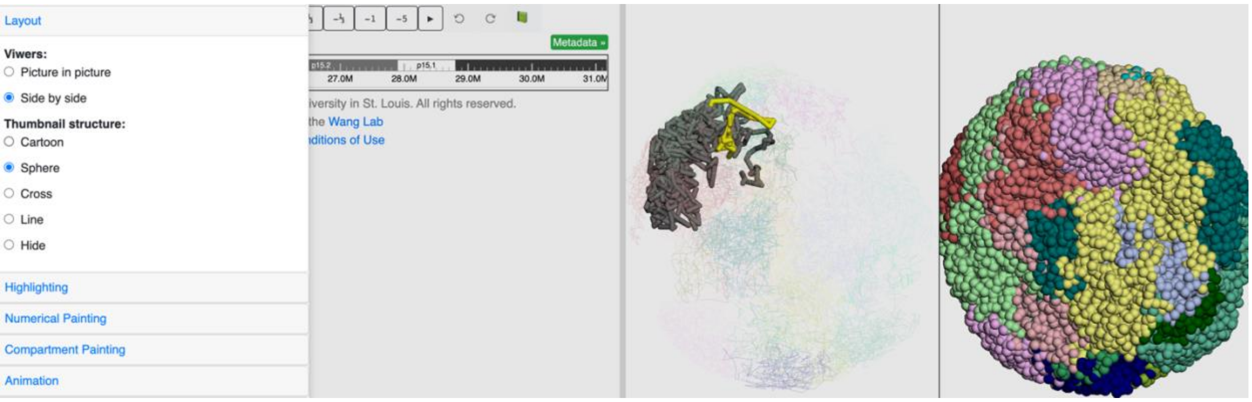

**Highlighting & labeling**

**Toggle browser region highlighting**

By default, the main viewer will highlight the structure of the sequence belonging to the linear browser region in yellow. The *Highlighting* section allows the user to click *Remove highlight* to turn off the highlighting.

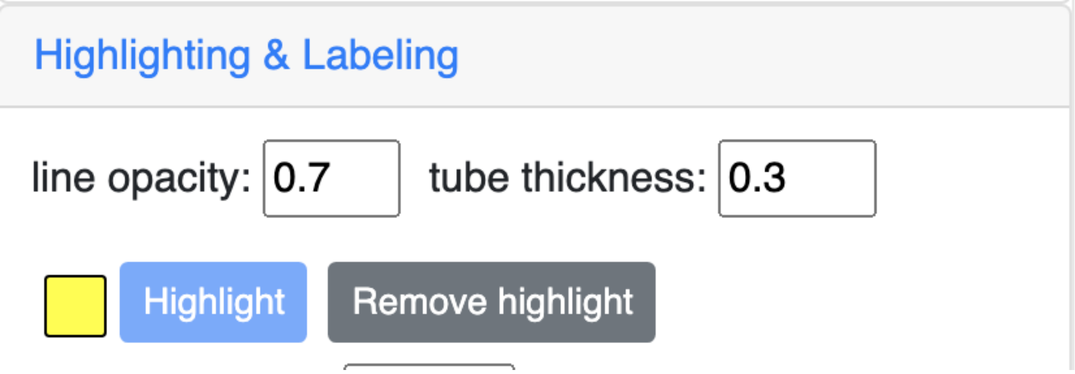

This is how it looks like when the highlighting is turned off:

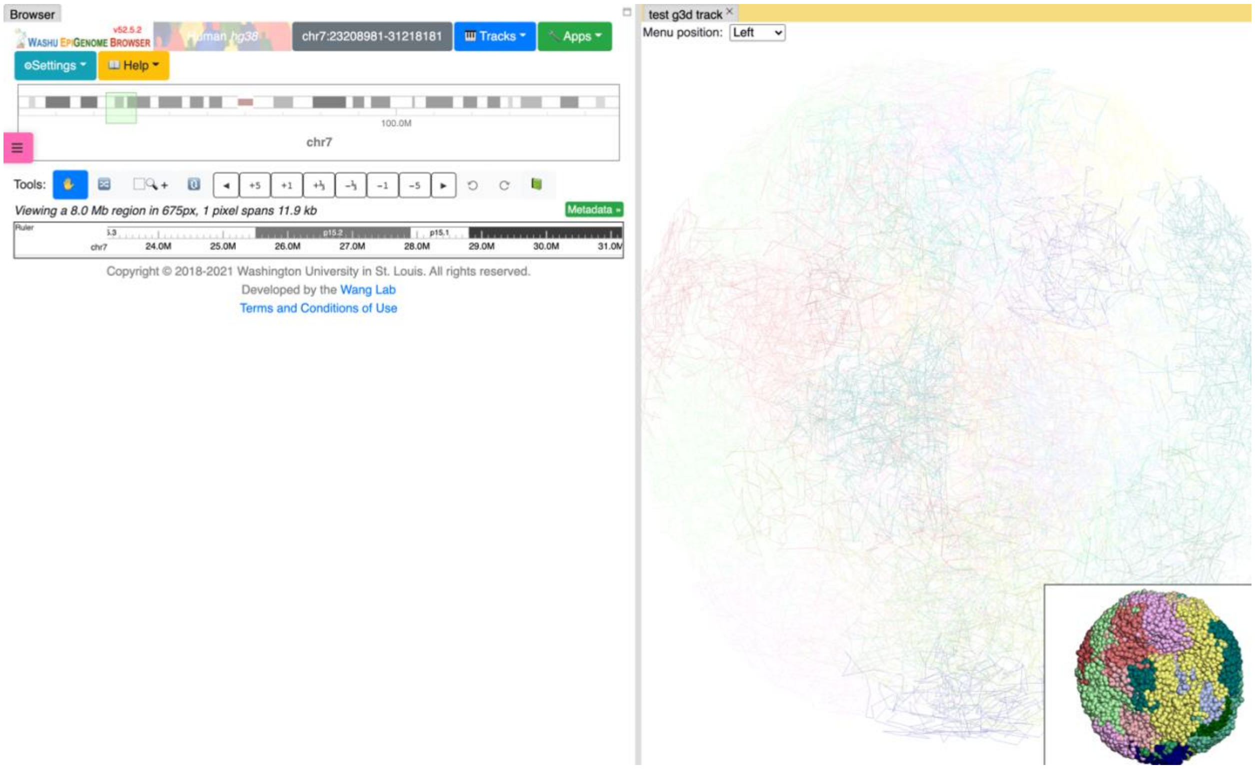

**Customize highlighting**

The highlighting color and tube thickness can also be customized. As shown below, we changed the color to purple and thickness to 1:

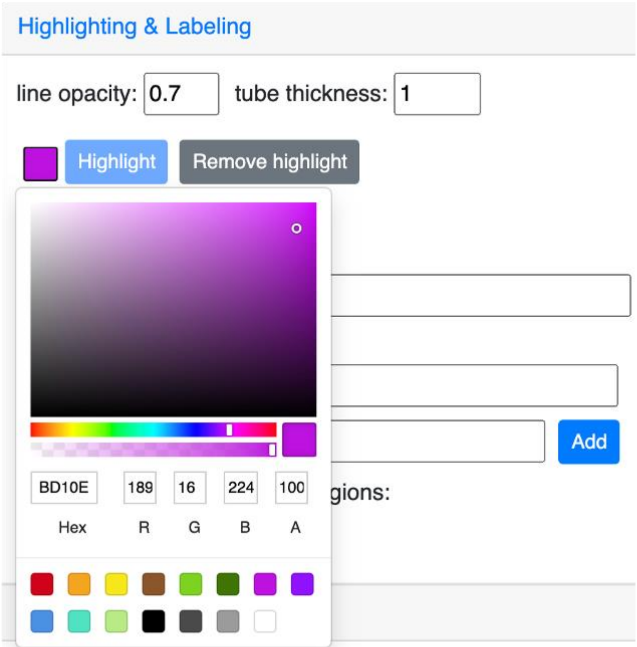

and this is the updated and view:

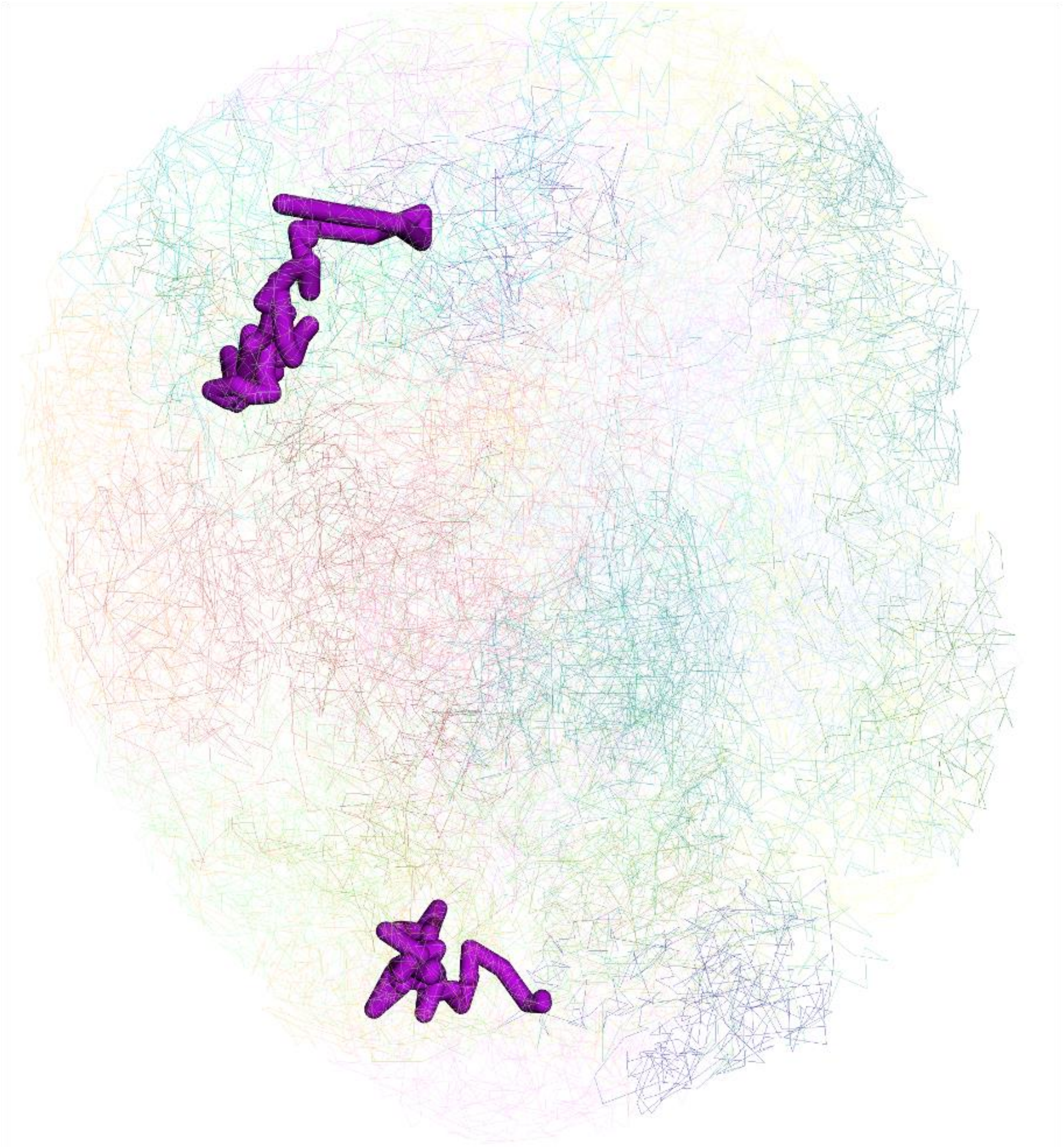

**Labeling by gene**

A gene symbol can be searched for labeling. To start, search any gene symbol and the menu will auto complete the search based on the user’s input.

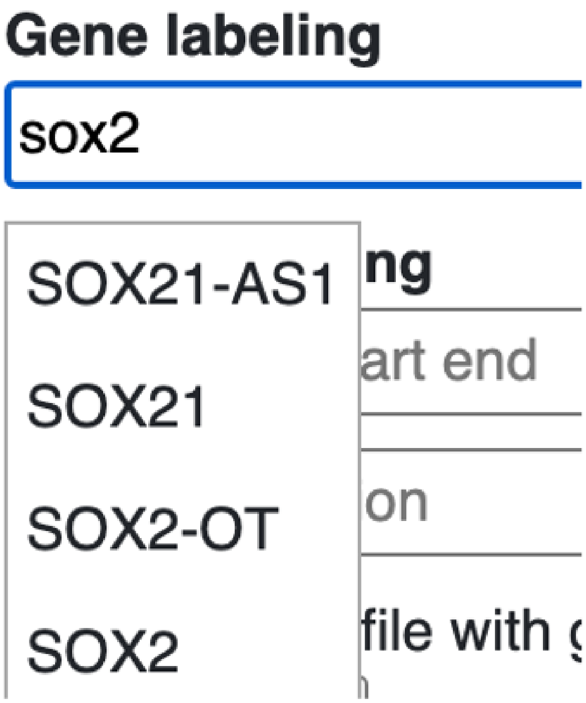

choose the correct isoform:

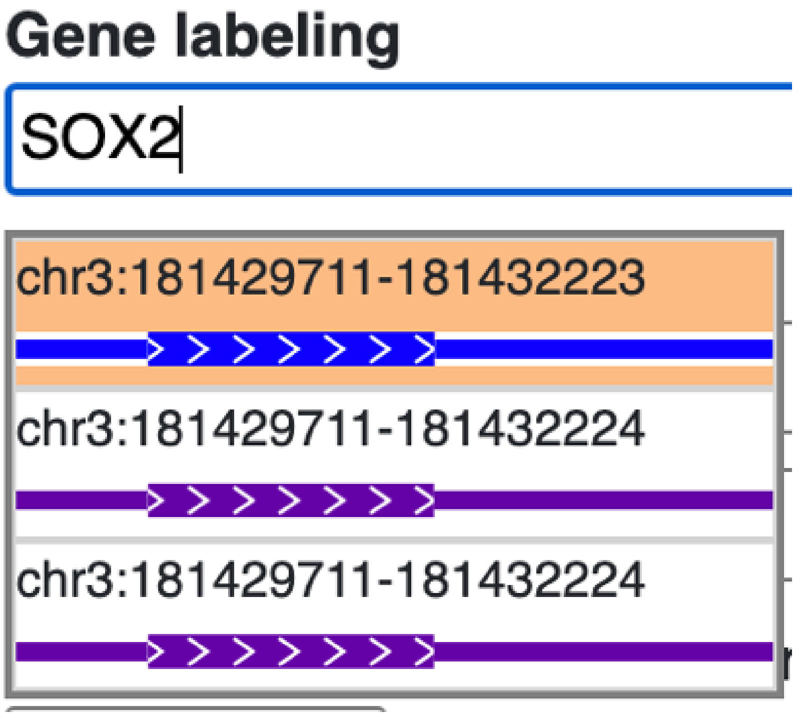

the gene will be added as a new label in the label list:

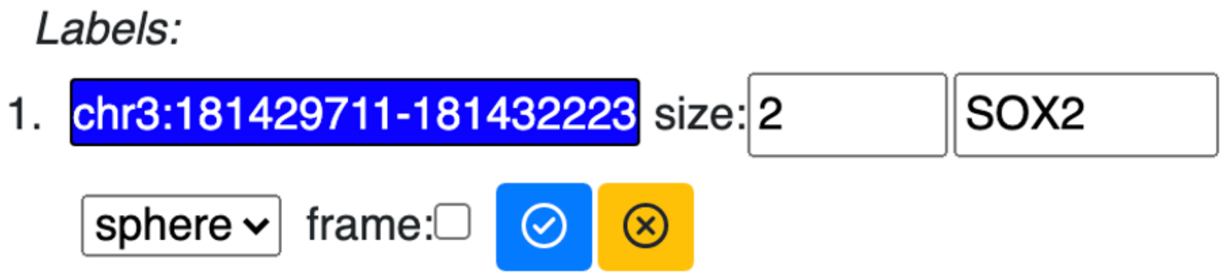

and shown in 3D view:

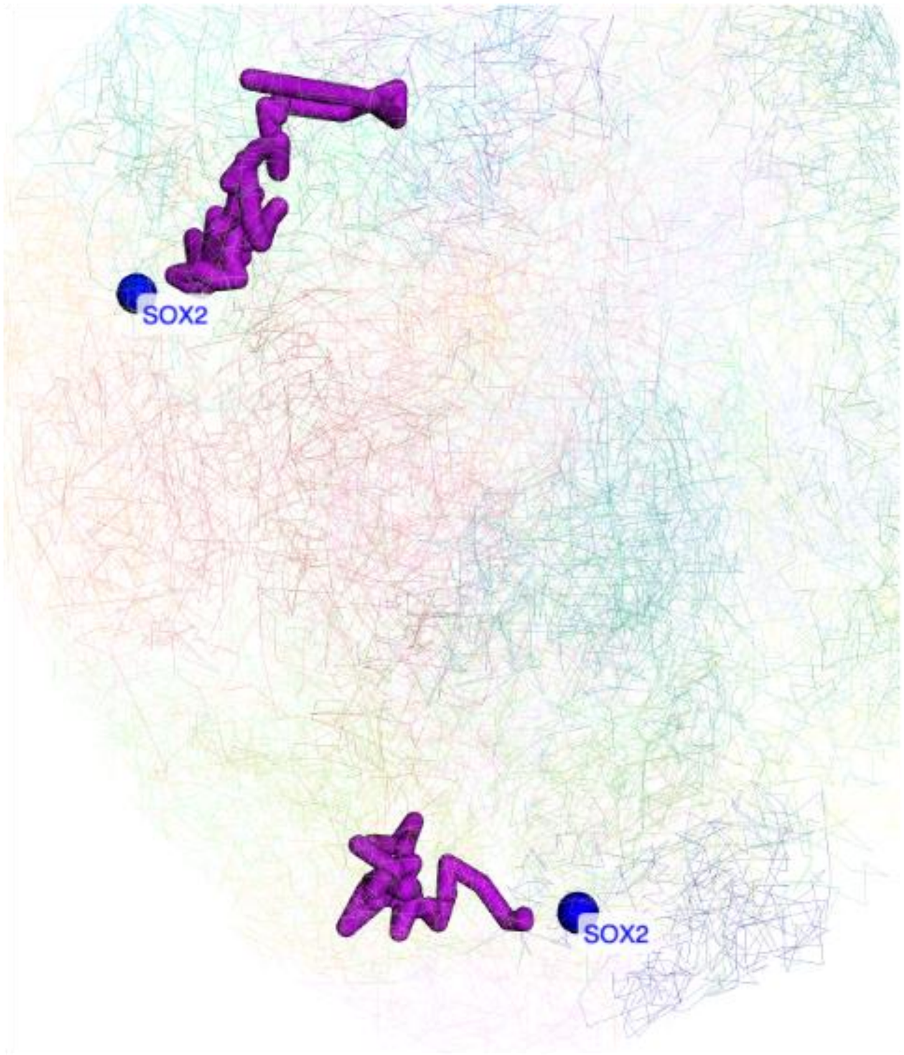

updated the display style of the label:

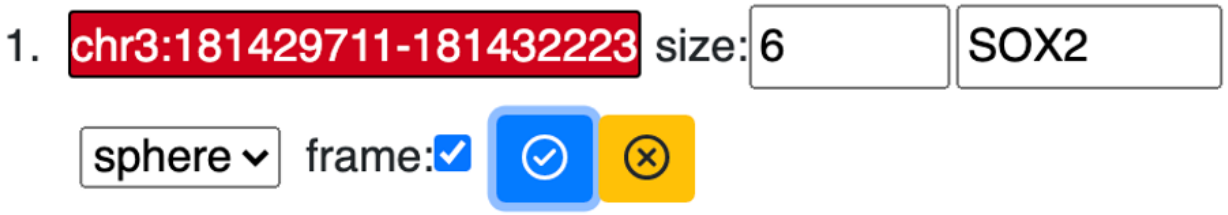

updated view of the label:

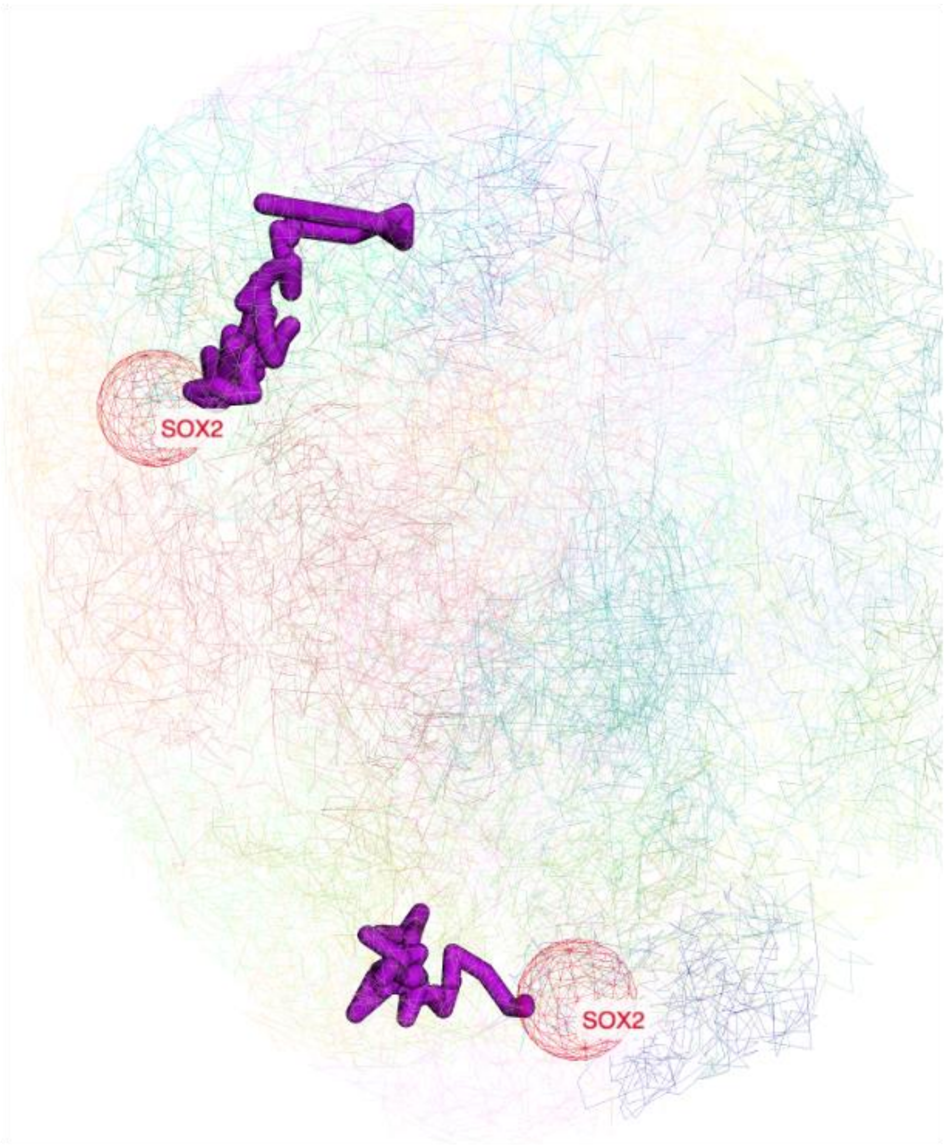

**Labeling by region**

User can also manually input a region for highlighting:

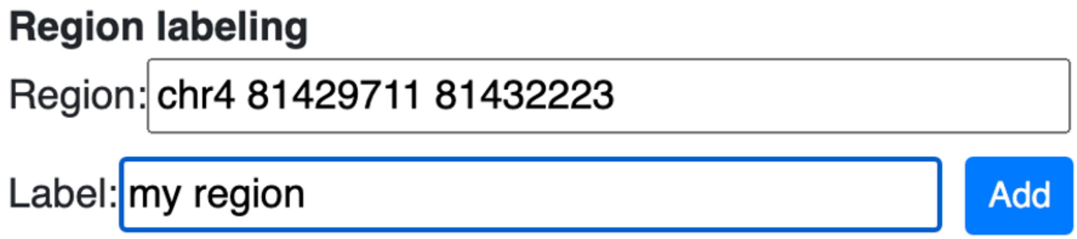

the added label name and region location will be updated in the menu control:

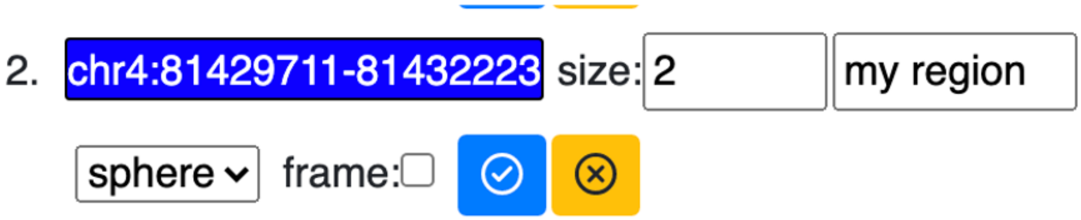

the view updated with the region location and corresponding label:

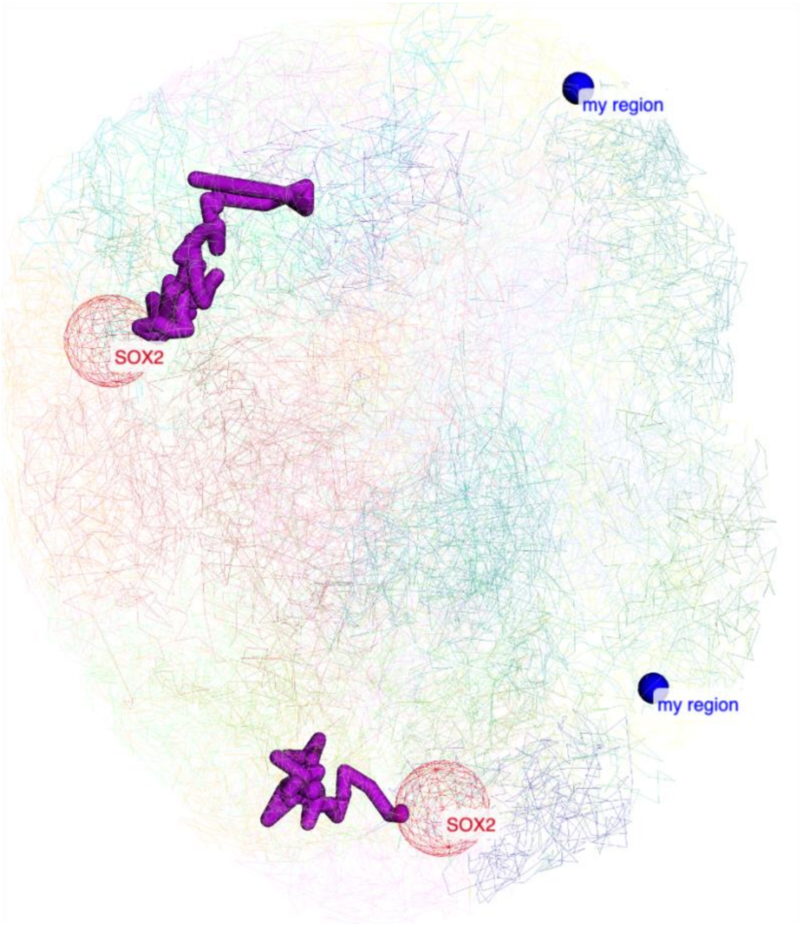

**Upload a file for labeling**

A text file containing a list of regions/gene symbols can also be uploaded for batch labeling, as shown below in the text file:

CYP4A22
chr10:96796528-96829254
CYP2A6
CYP3A4
chr1:47223509-47276522
CYP1A2

upload this file:

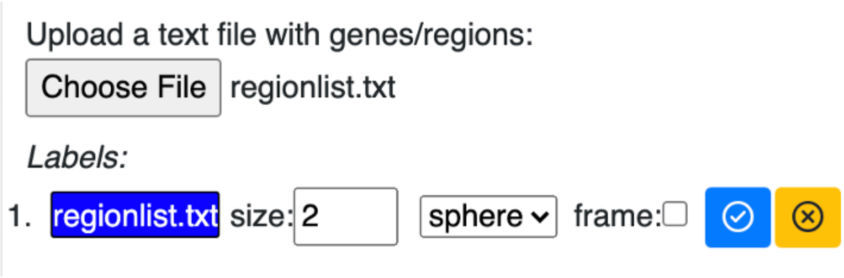

regions in the file are all labeled:

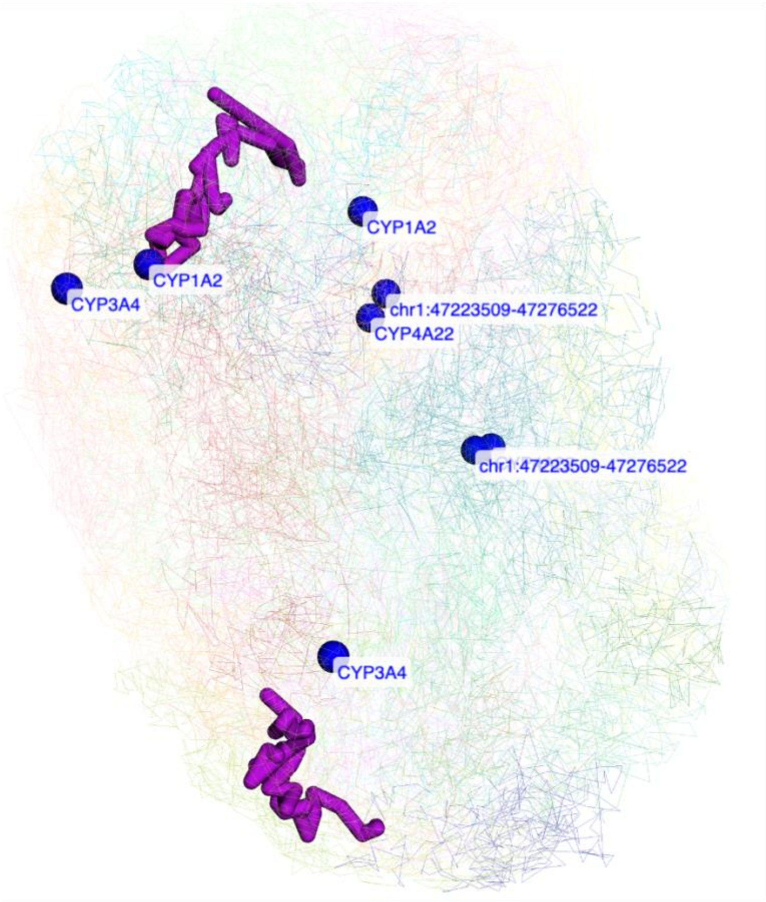

**Pointing using arrows**

In addition to using shapes for labels, arrows can also be used to point to the selected region. Choose label style as arrow:

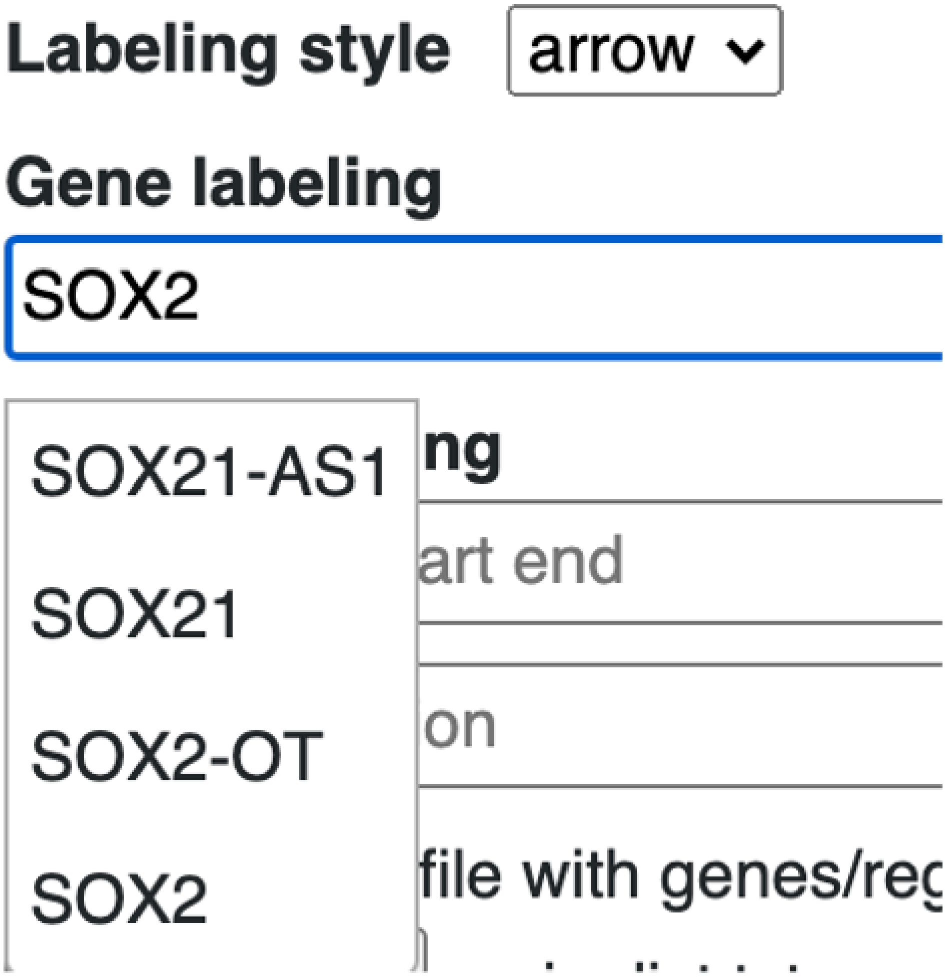

use either the gene search or region labeling:

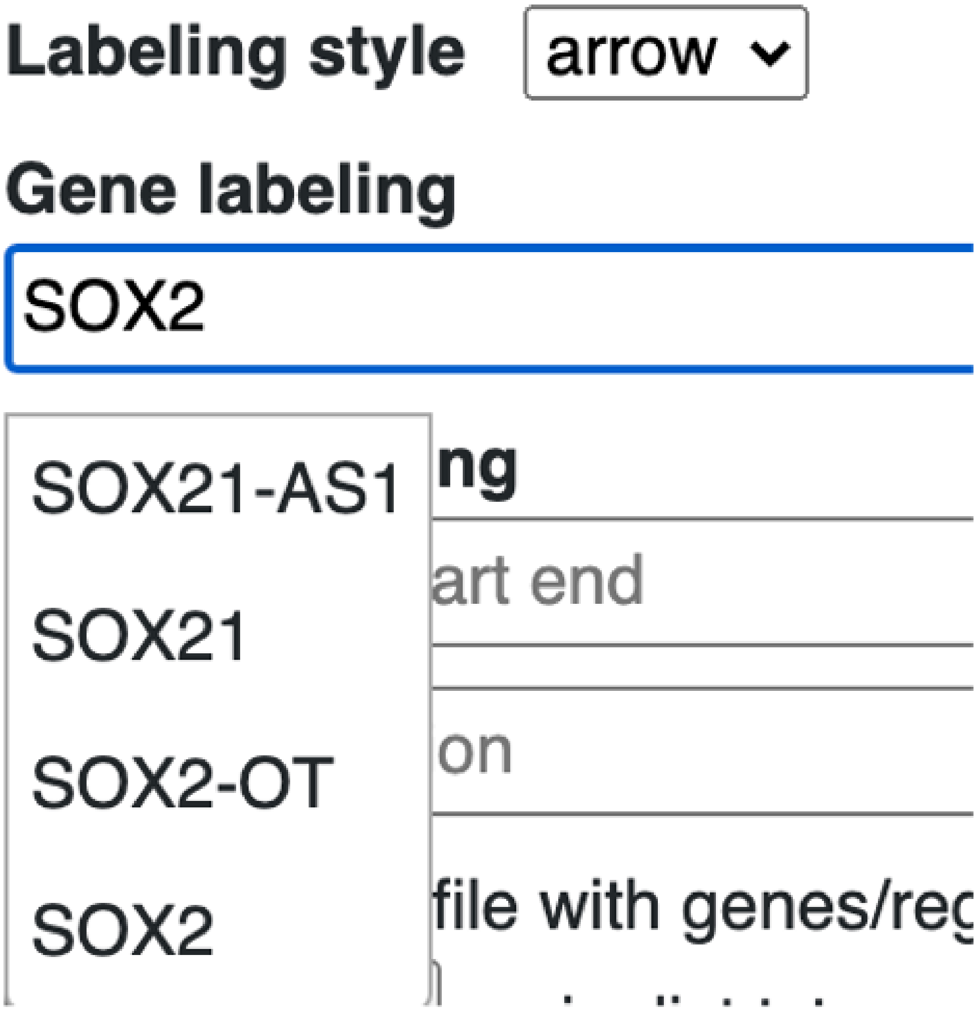

the new label will be displayed under the arrow list:

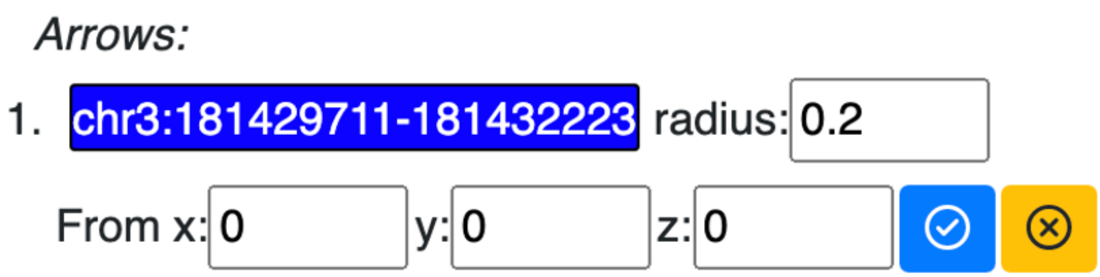

and displayed in the 3D viewer:

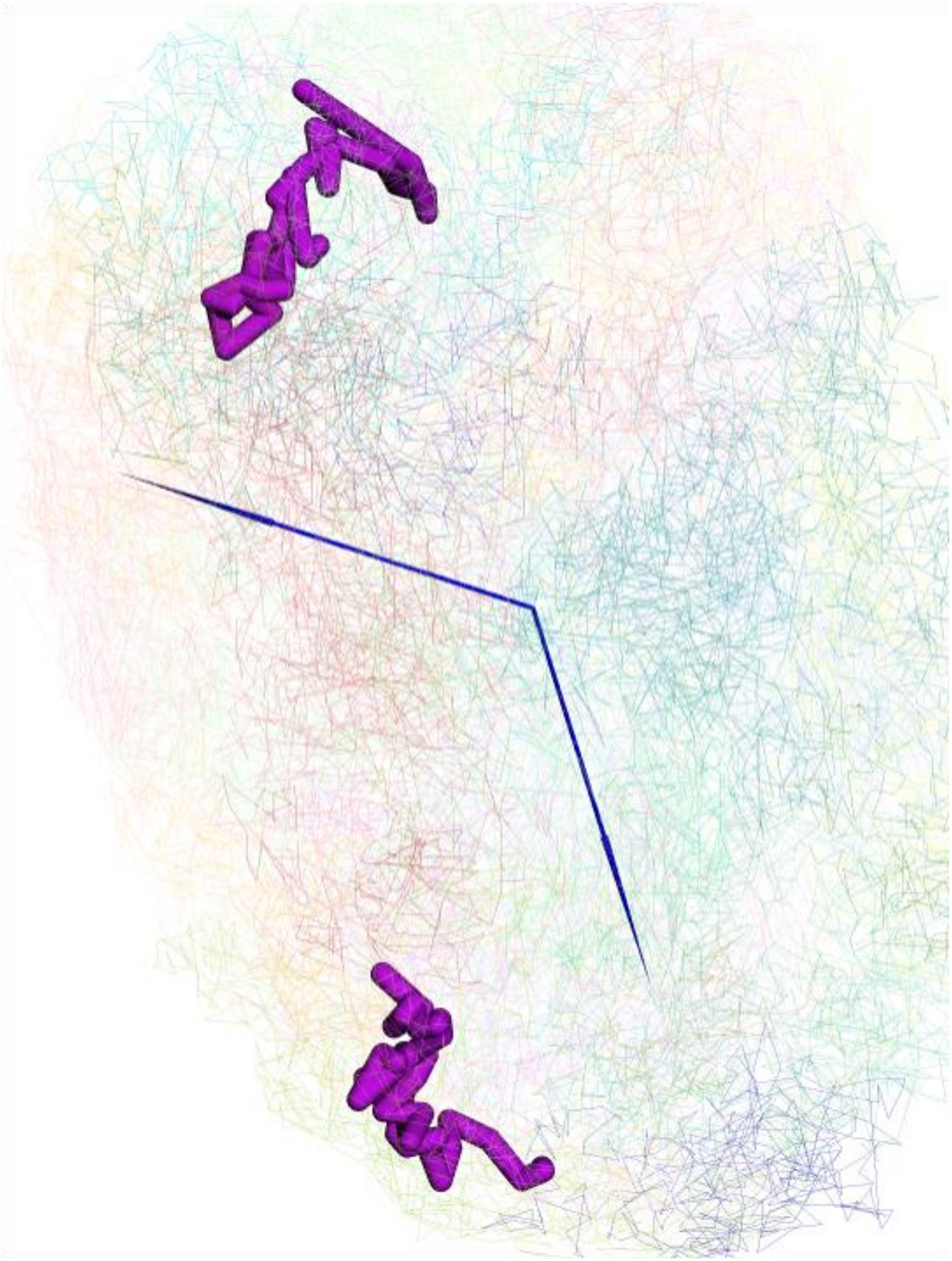

configure the style of the arrow:

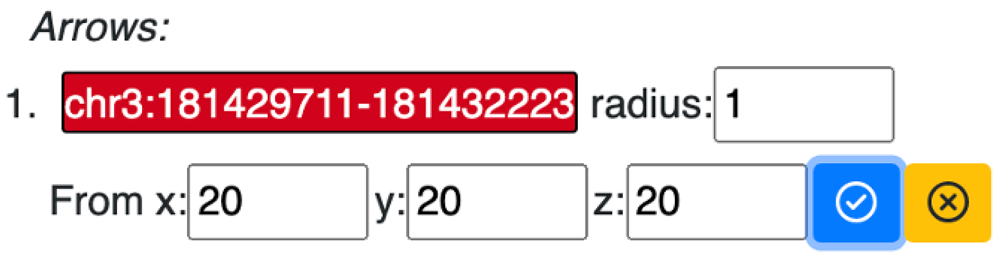

updated arrow style in the viewer:

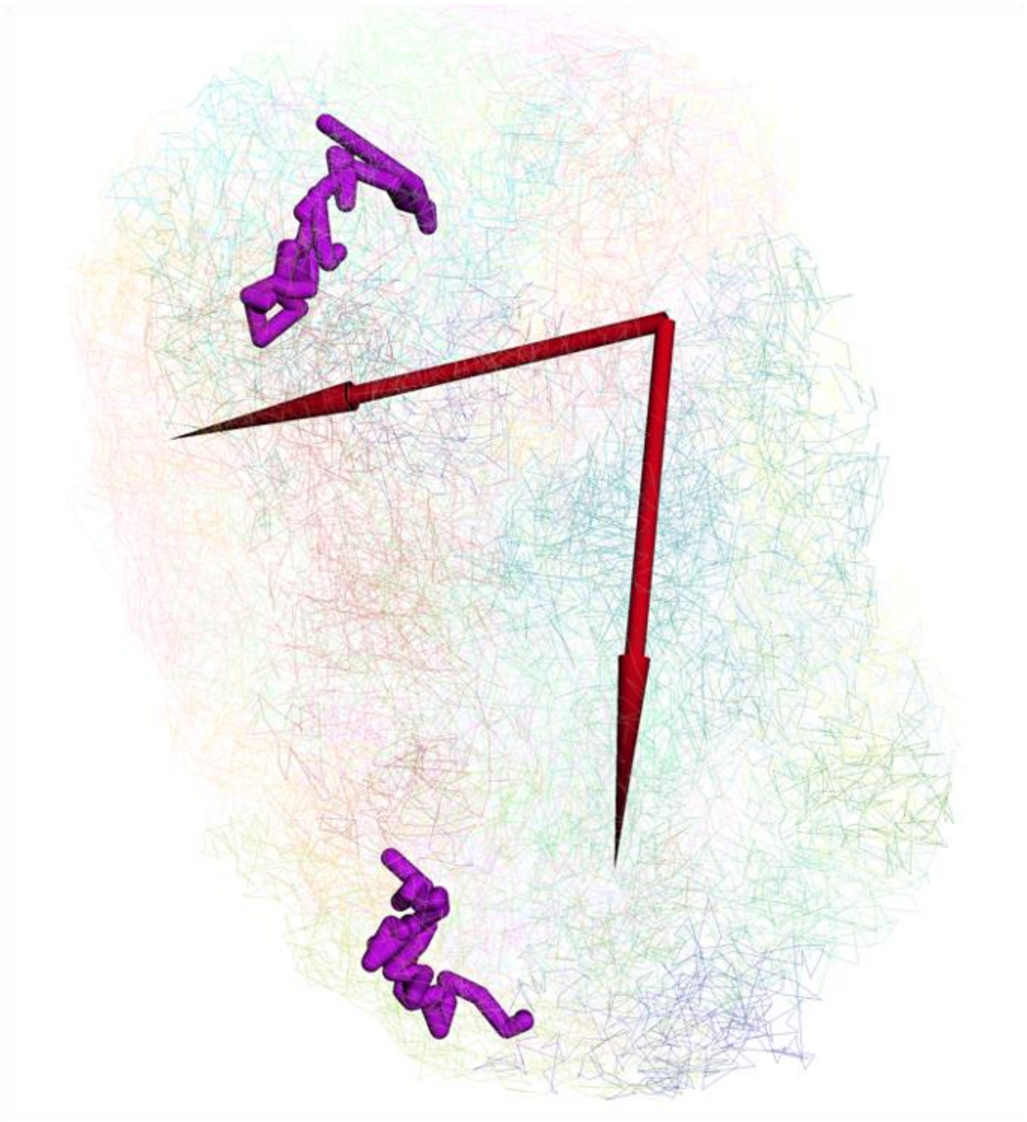

**Interactivity between viewers**

For certain track types, like gene annotation and Hi-C tracks, users can choose to display gene or Hi-C anchors on the 3D structure directly. As shown below, the gene tooltip has a

Show in 3D button, click this button to add this gene to the label list and highlight it in 3D view:

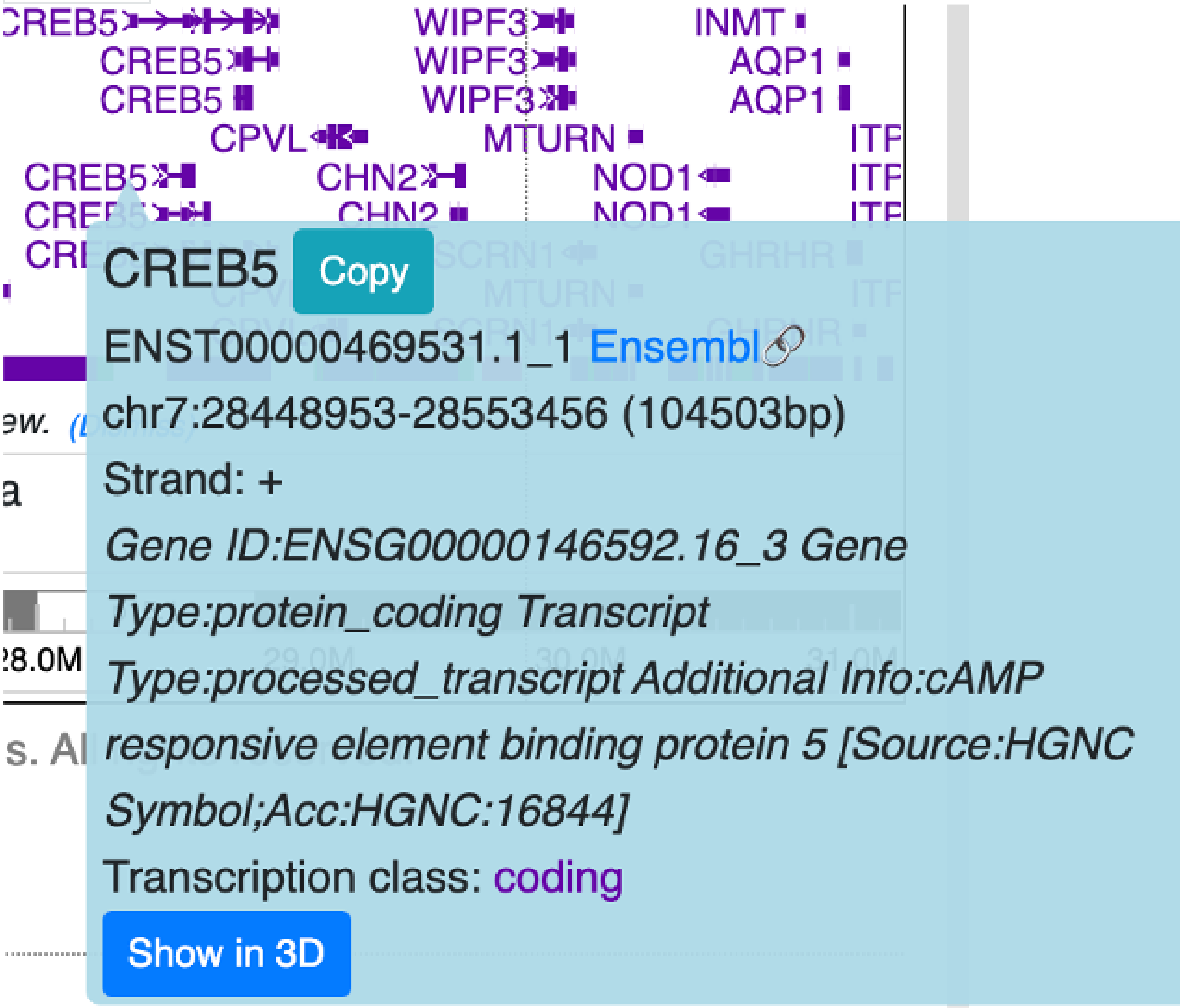

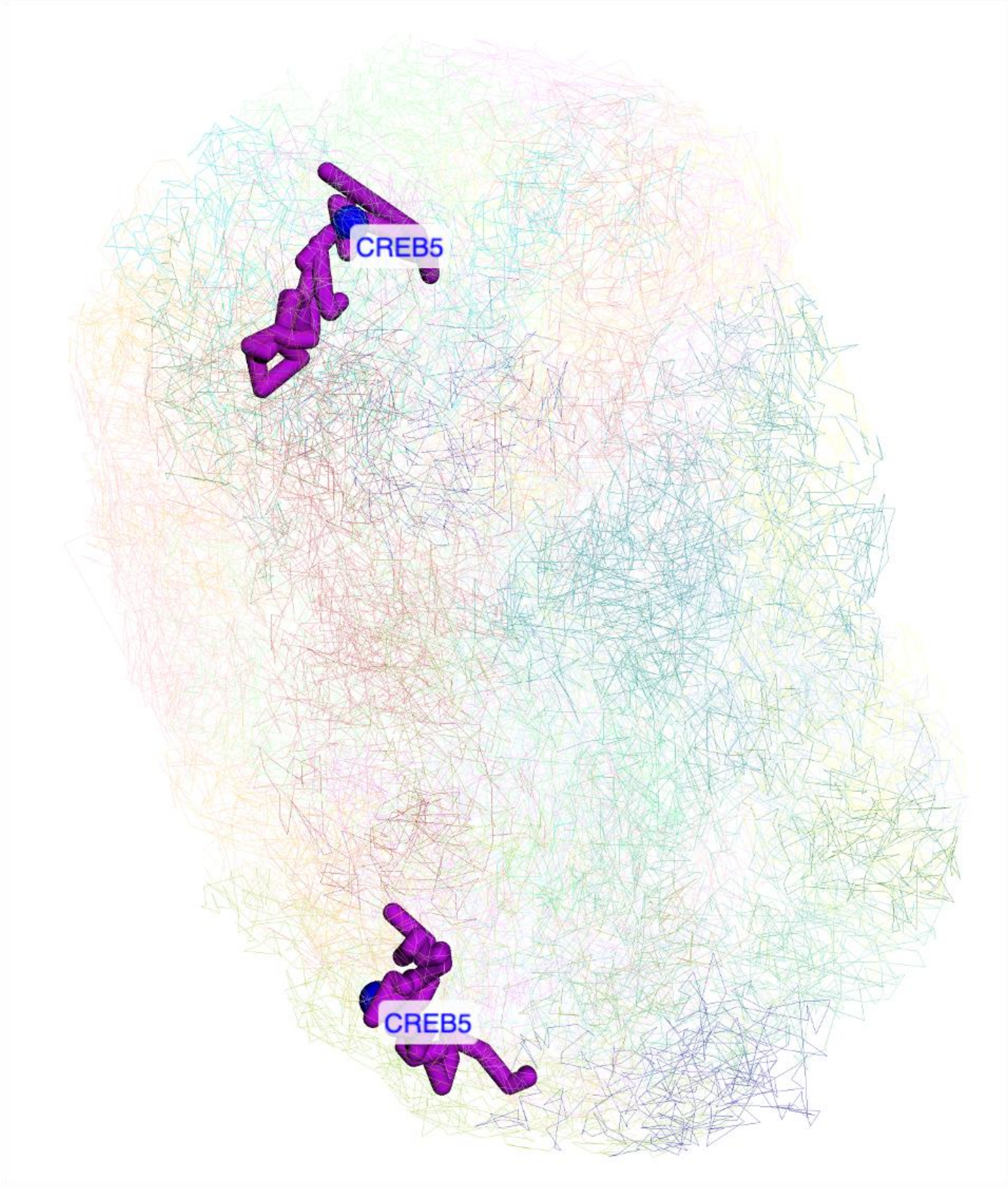

Clicking any square on a Hi-C track will also bring the Show in 3D button, click it to add both of the contact anchors to the arrow list:

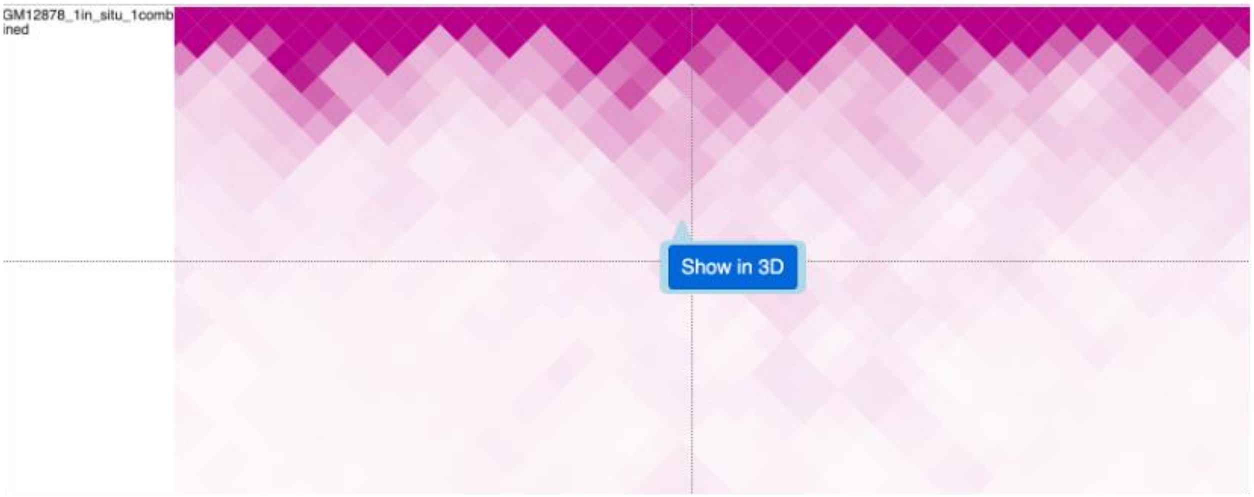

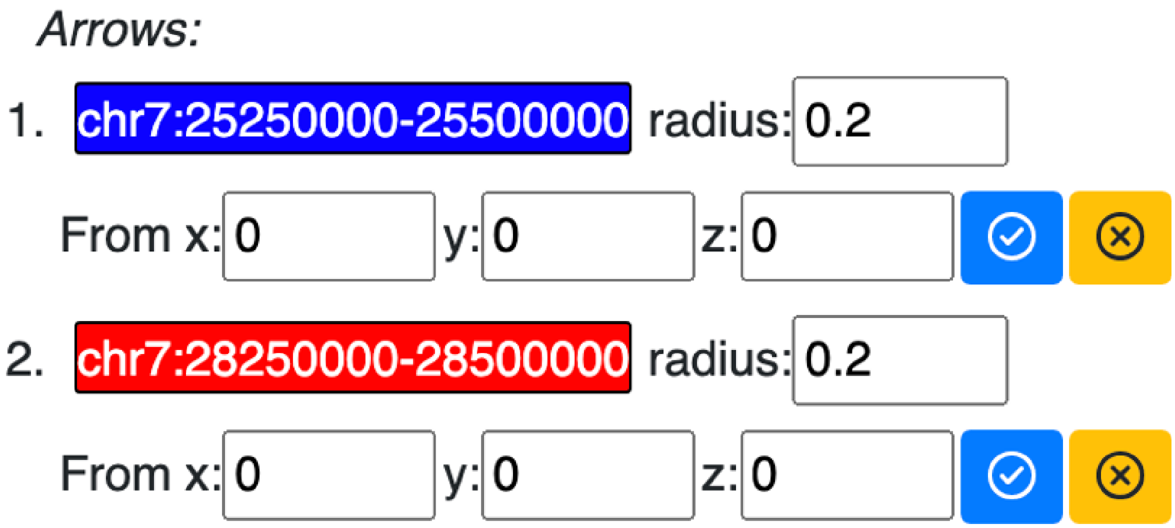

arrows pointing to both anchors will be displayed in the 3D view. Since both the paternal and maternal models are visible in this structure, there are 4 arrows displayed, each representing the two anchors per model:

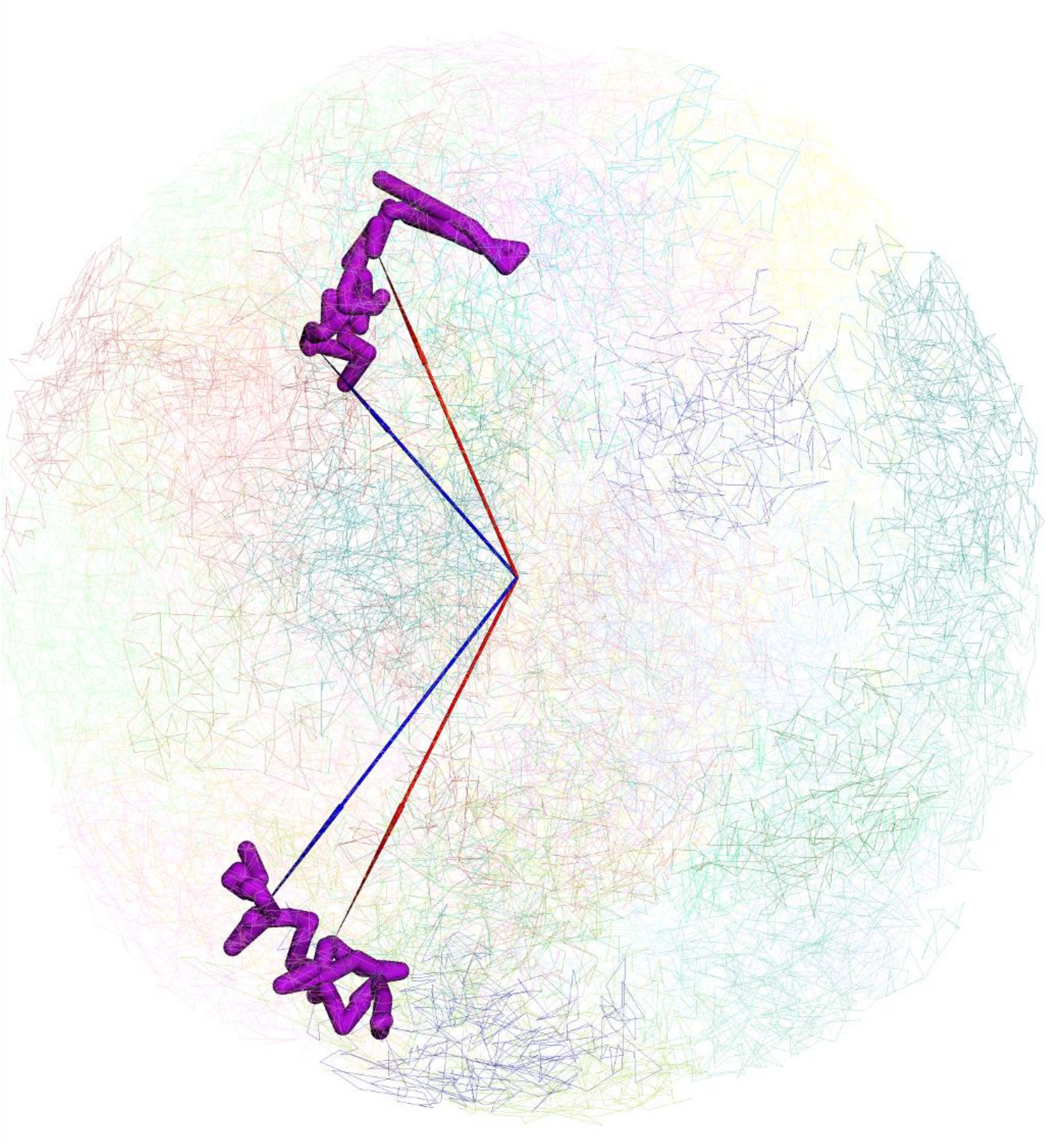

**Numerical painting**

**Numerical painting with bigwig data**

The numerical track in bigWig format can be used to paint the 3D structure. The *Use loaded tracks* check option allows users to load either bigWig tracks already loaded on the browser or submit another bigWig track with the file URL.

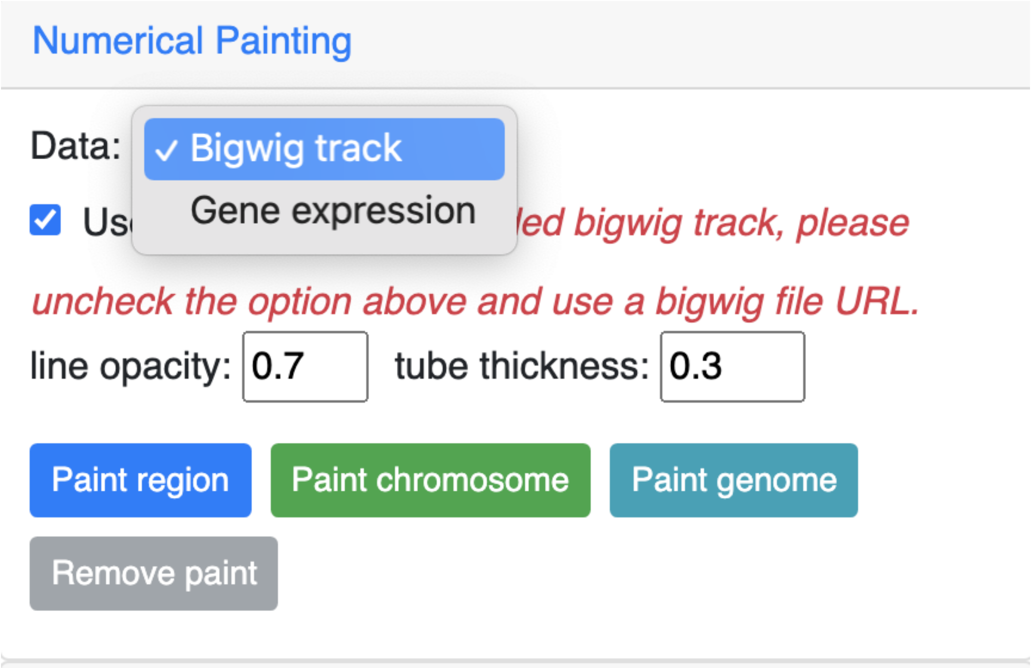

If *Use loaded tracks* is unchecked, the user submits the bigWig URL input:

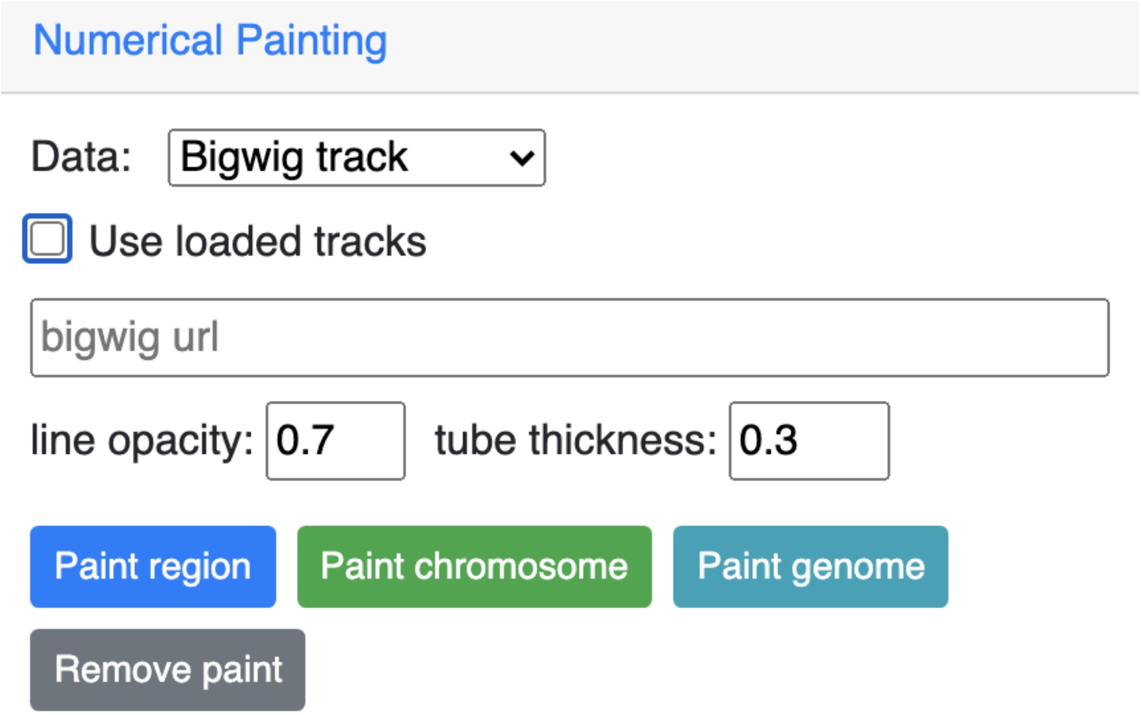

Here, we are using the GC percentage data of *hg19* genome as example, add the *GC Percent* track from *Annotation Tracks*:

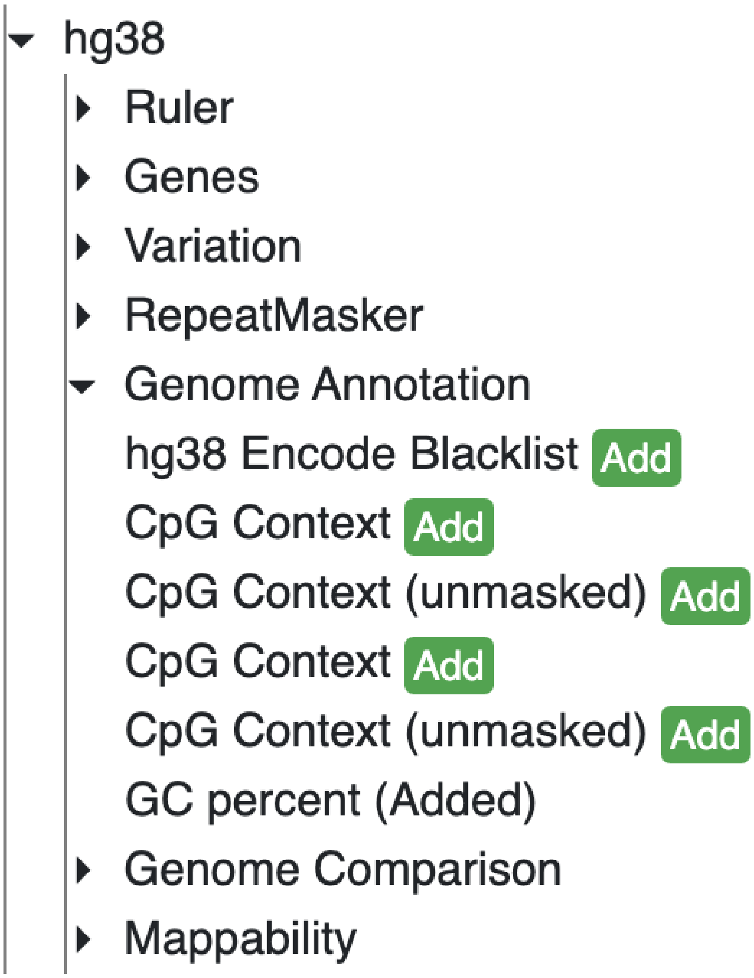

After the GC Percent track is added. choose the track from the dropdown menu:

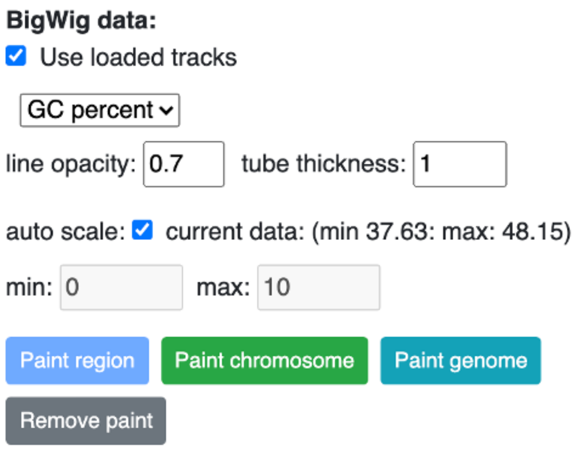

Click *Paint region* button:

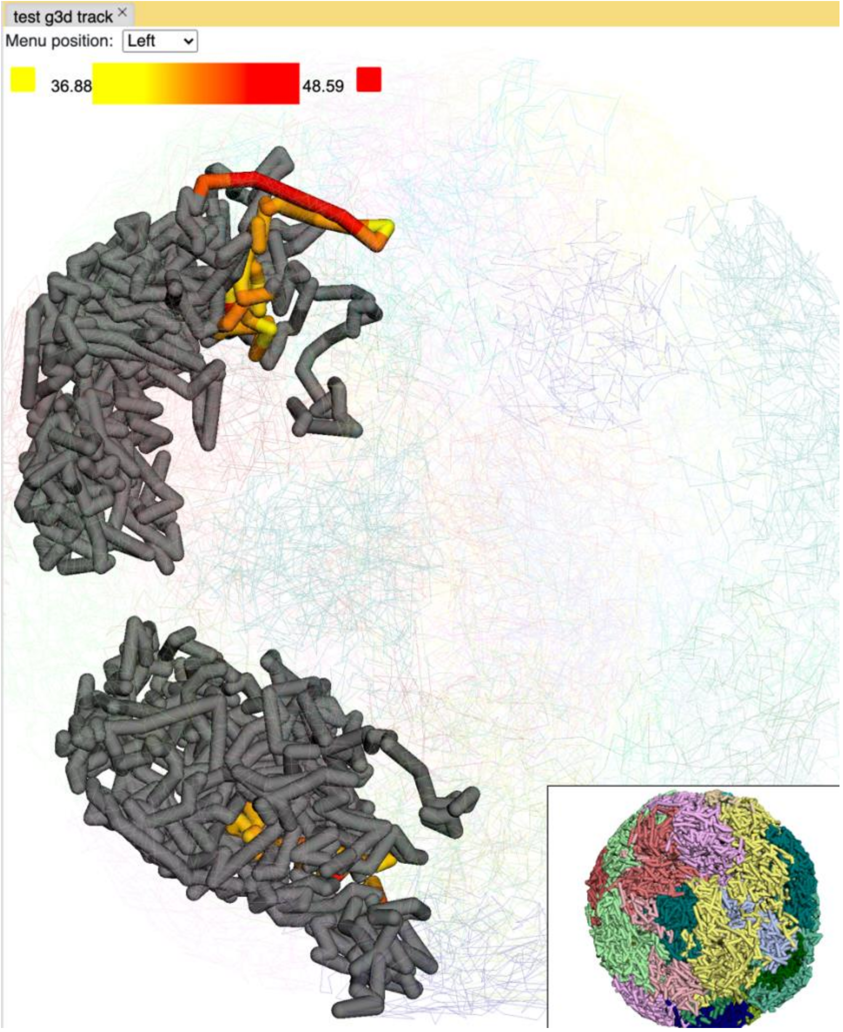

you can also paint the whole chromosome by click the *Paint chromosome* button:

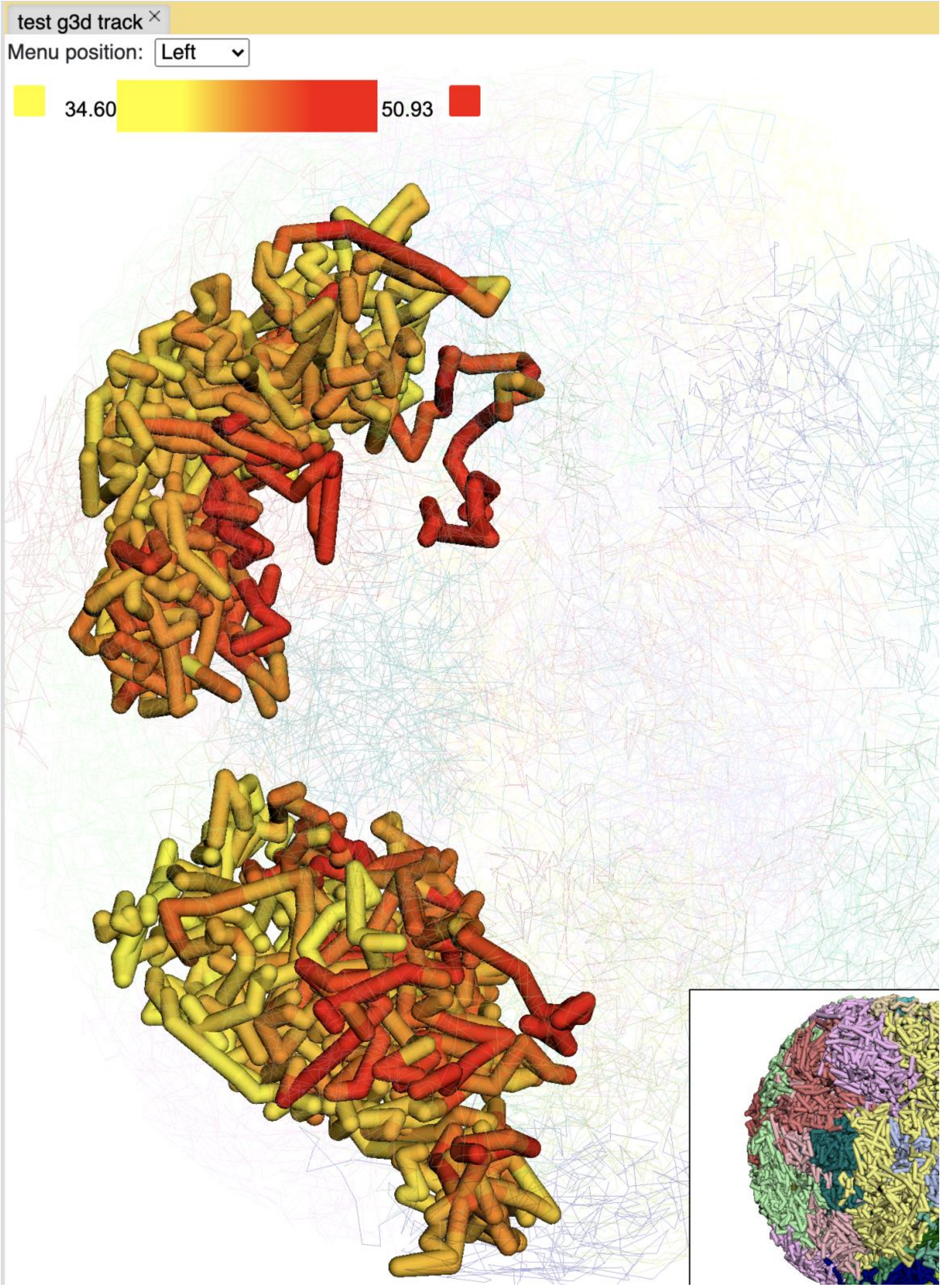

Click the color box on the color legend to bring up a color palette to select colors:

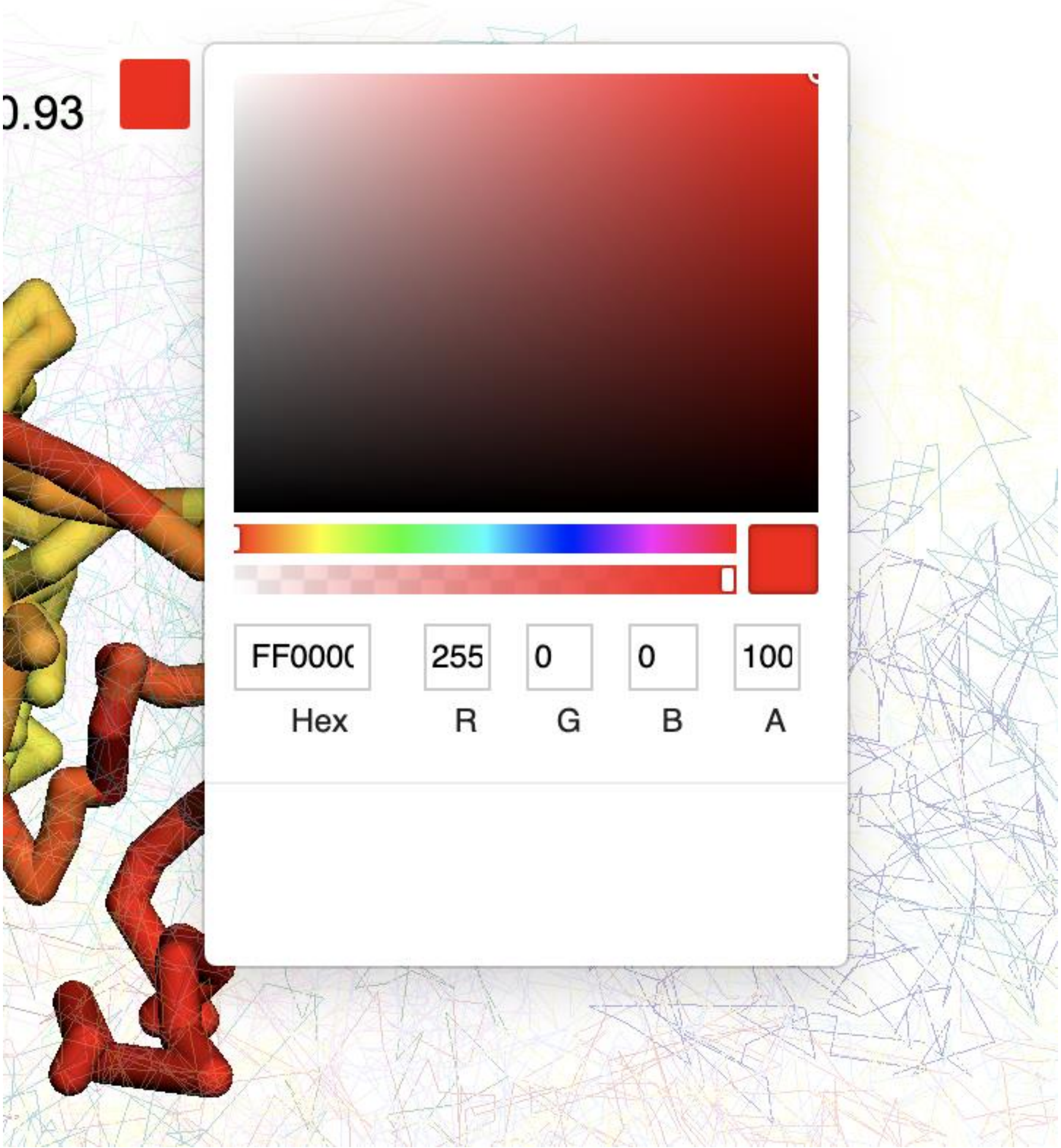

Choosing a different color will automatically reload the structure with the color chosen:

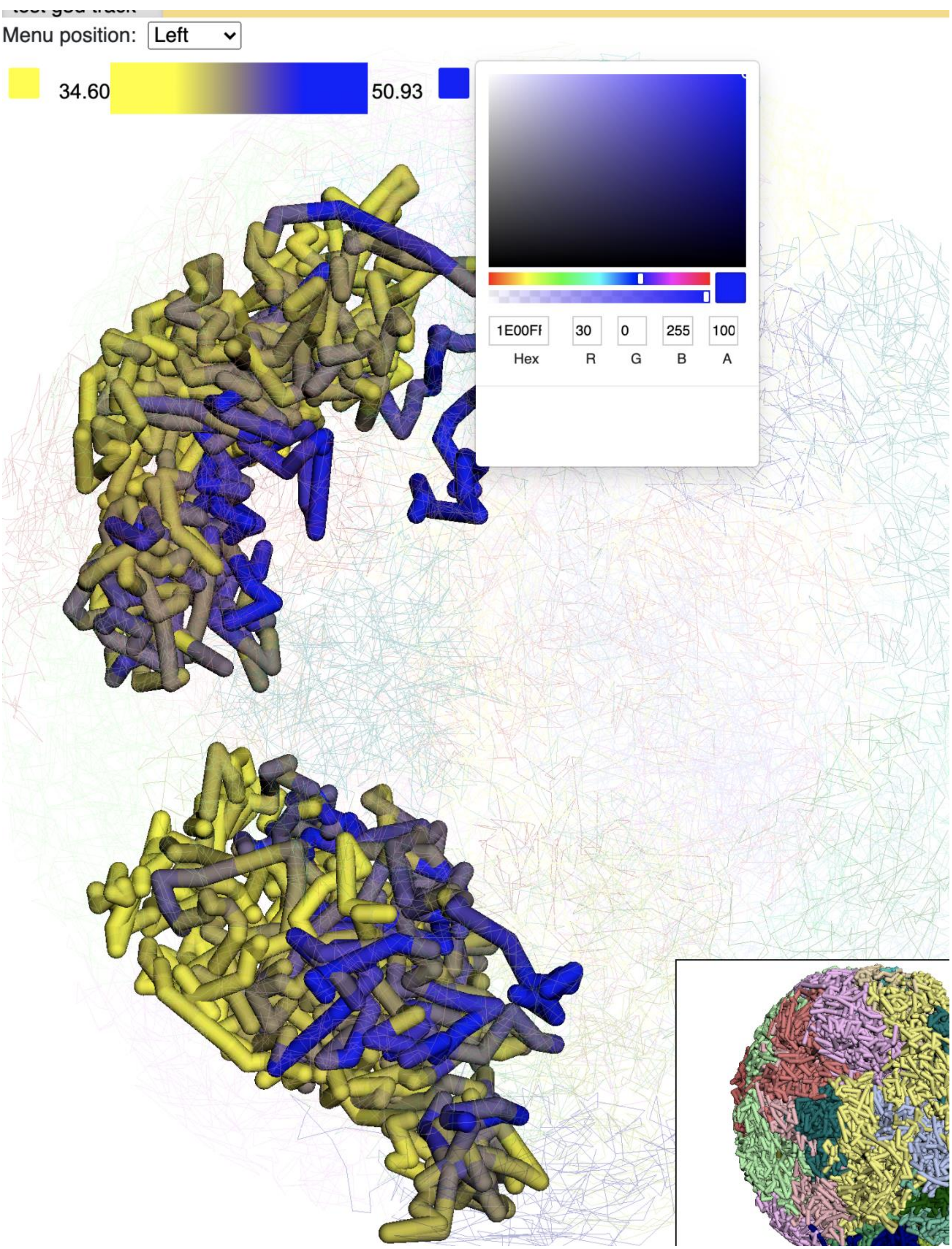

Menu position: Left
34.60
50.93
1E00FF
30
0
255
100
Hex
R
G
B
A

To see the whole genome painted, click *Paint genome*:

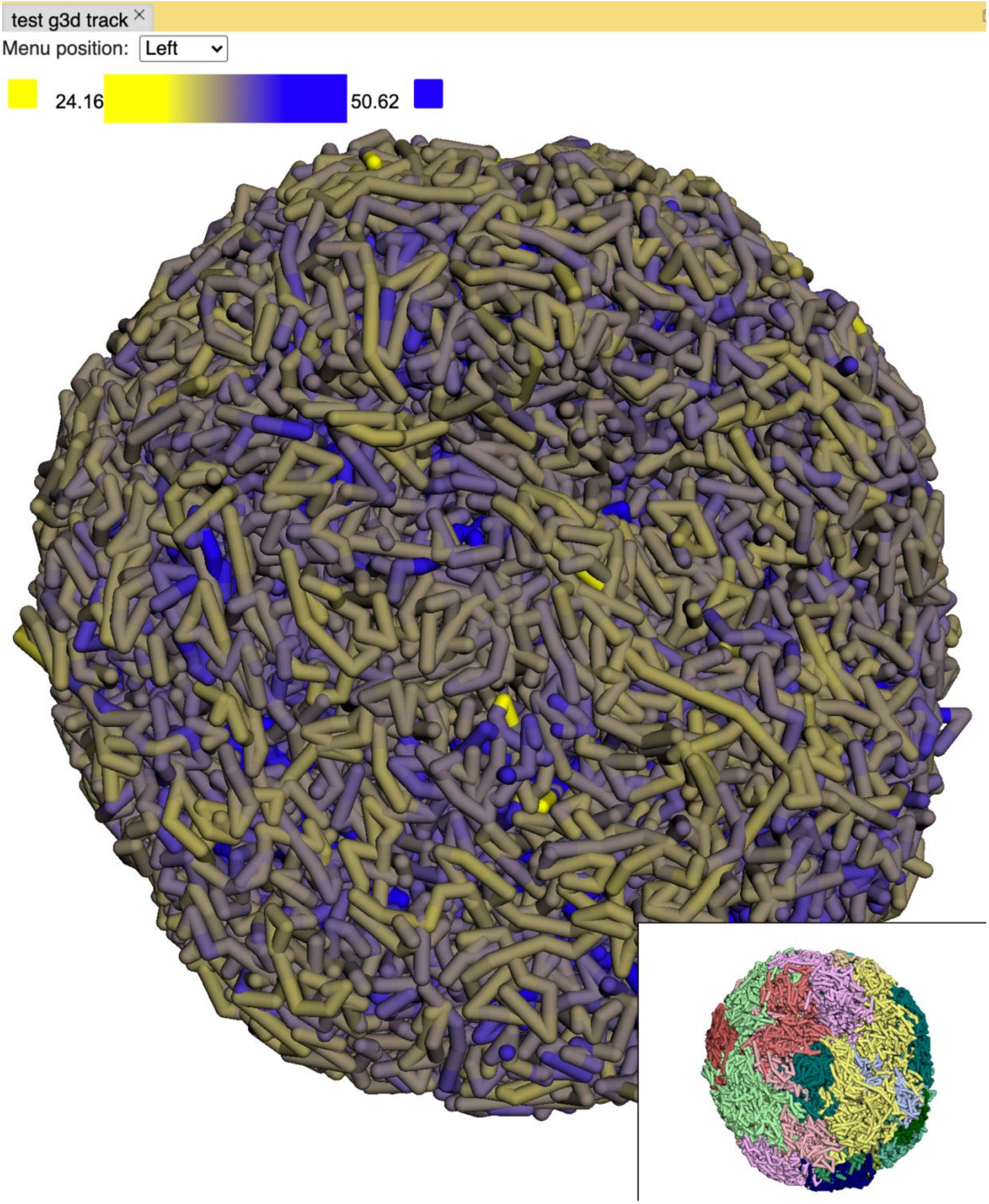

Click the *Remove paint* button to remove the painting.

**Numerical painting with gene expression data**

For painting with gene expression data, the data needs to be organized in the following format:

| | | | | |
|---|---|---|---|---|
| chr3 | 168903366 | 168921996 | ENSG00000242268.2 | 2.40146671319 |
| chr18 | 46756487 | 46764408 | ENSG00000270112.3 | 0.0287250976522 |
| chr3 | 11900011 | 11901245 | ENSG00000225275.4 | 0.0 |
| chr15 | 41921417 | 41928883 | ENSG00000259883.1 | 0.305029986379 |
| chr13 | 98949719 | 98950447 | ENSG00000231981.3 | 0.0806509326125 |
| chrX | 152682810 | 152683842 | ENSG00000269475.2 | 0.0 |
| chr12 | 44880868 | 44880969 | ENSG00000201788.1 | 0.0 |
| chr17 | 57092145 | 57096425 | ENSG00000263089.1 | 0.295277363304 |

This is a 5 column bed format file (chromosome, start, end, gene id or symbol, gene expression value) and can be FPKM, RPKM or whatever type of value you want to plot.

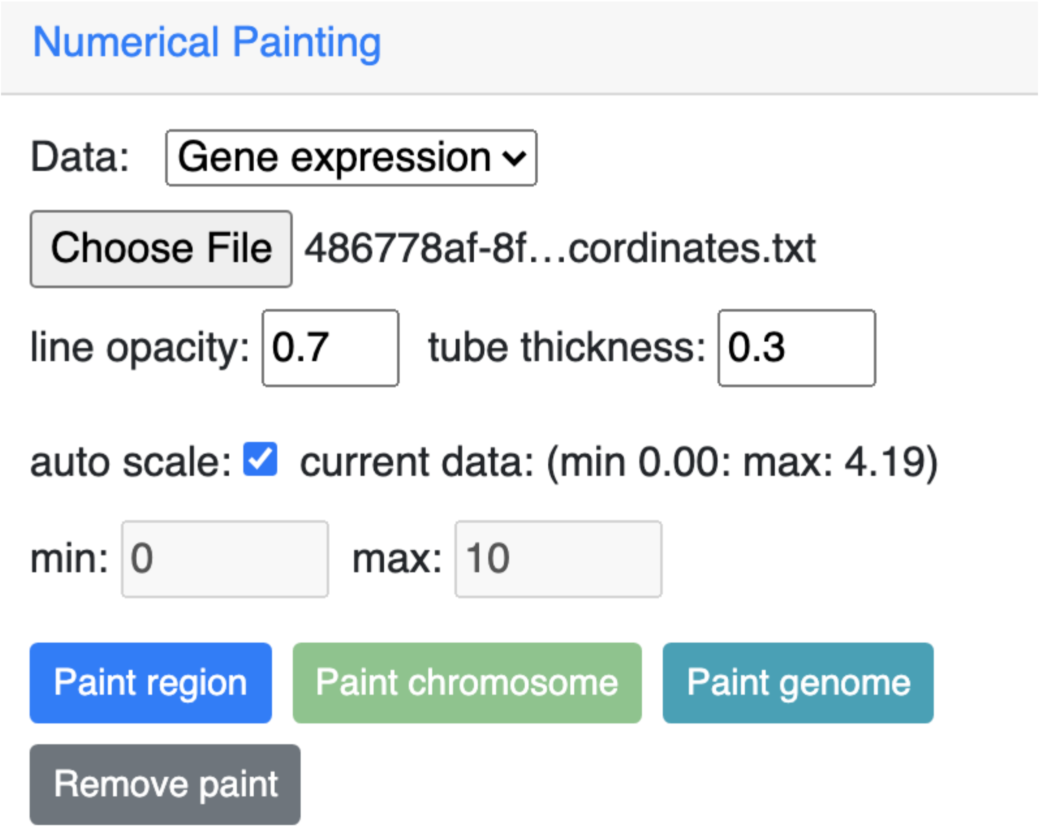

Choose *Gene expression* from the dropdown menu, then upload your file and click one of the paint buttons.

This is the view after painting with the expression data. The color and scale can be customized as described before:

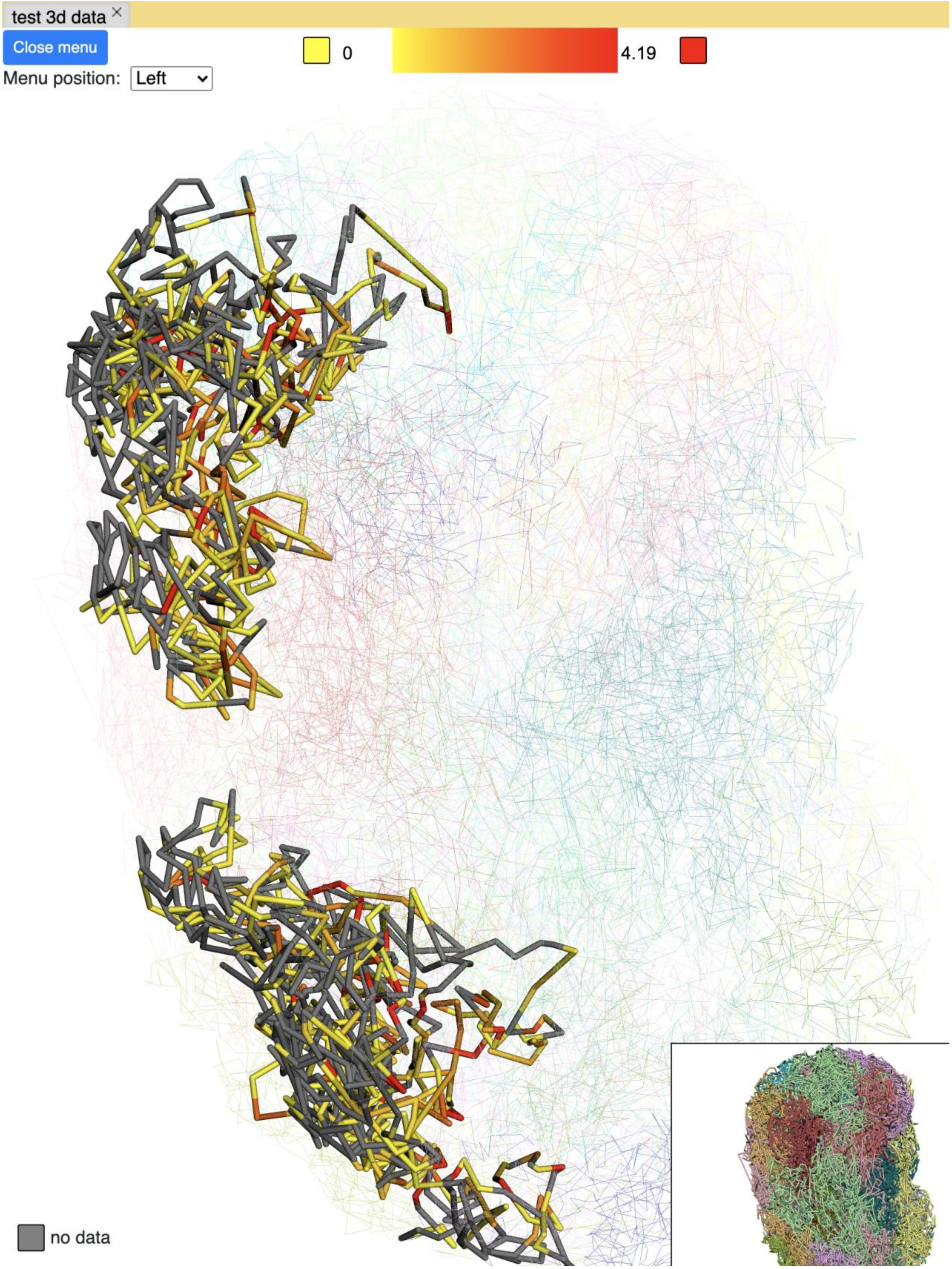
test 3d data
Close menu
Menu position: Left
0
4.19
no data

**Annotation painting**

**Supported file formats for 3D annotation painting**

**cytoband**

For *cytoband* there is no need to upload a file, the cytoband data will be read from the current loaded genome data.

**refGene**

The standard *refGene* format from UCSC can be used for painting gene positions on the 3D model:

```
2085	NR_046630	chr3	+	196666747	196669405	196669405	196669405	3	196666747,196667841,196669263,	196666995,196668013,196669405,	0	NCBP2-AS1	unk	unk	-1,-1,-1,
2051	NR_046598	chr3	+	192232810	192234362	192234362	192234362	2	192232810,192234269,	192233297,192234362,	0	FGF12-AS2	unk	unk	-1,-1,
1312	NR_046514	chr13	+	95364969	95368199	95368199	95368199	2	95364969,95365891,	95365647,95368199,	0	SOX21-AS1	unk	unk	-1,-1,
585	NR_106918	chr1	-	17368	17436	17436	17436	1	17368,	17436,	0	MIR6859-1	unk	unk	-1,
585	NR_107062	chr1	-	17368	17436	17436	17436	1	17368,	17436,	0	MIR6859-2	unk	unk	-1,
```

**bed 9 columns**

A bed file with the 9th column as RGB values can be used as well, for example the chromHMM from Roadmap project is shown below:

```
chr10	0	94800	15_Quies	0	.	0	94800	255,255,255
chr10	94800	95600	9_Het	0	.	94800	95600	138,145,208
chr10	95600	102200	15_Quies	0	.	95600	102200	255,255,255
chr10	102200	104400	9_Het	0	.	102200	104400	138,145,208
chr10	104400	110000	15_Quies	0	.	104400	110000	255,255,255
chr10	110000	111200	9_Het	0	.	110000	111200	138,145,208
```

**bed 4 columns**

To make things simple, a 4 column bed format is also supported, where the 4th column contains the color value:

| | | | |
|---|---|---|---|
| chr11 | 108280000 | 109080000 | *#ff0100* |
| chr11 | 109080000 | 109480000 | *#0000ff* |
| chr11 | 109720000 | 110160000 | *#018100* |
| chr11 | 110200000 | 111400000 | *#0064fb* |
| chr11 | 111400000 | 112640000 | *#ef8c0a* |
| chr11 | 112640000 | 113480000 | *#7f007f* |
| chr11 | 113520000 | 114520000 | *#520000* |
| chr11 | 114520000 | 114880000 | *#39ae00* |

**4DN compartment data**

A file with a compartment call table  can also be used to paint the 3D structure. We provide the compartment call data for HeLaS3 (4DNFIL65C8ZI) from the 4DN data portal. The file is relatively small ~about 1MB in size. The file can either be in raw text format (example text) or in compressed gzip format (example gzipped text) for upload.

The 4DN compartment data looks like:

| chrom | start | end | gene_count | gene_coverage | E1 | E2 | E3 |
|---|---|---|---|---|---|---|---|
| chr1 | 0 | 100000 | 595 | 0.8812700000000001 | | | |
| chr1 | 100000 | 200000 | 952 | 1.0 | | | |
| chr1 | 200000 | 300000 | 159 | 0.09797 | | | |
| chr1 | 300000 | 400000 | 132 | 0.05368 | | | |
| chr1 | 400000 | 500000 | 471 | 0.24454 | | | |
| chr1 | 500000 | 600000 | 390 | 0.15467999999999998 | | | |
| chr1 | 600000 | 700000 | 229 | 0.05782999999999999 | | | |

**Rao et.al compartment data**

Rao et.al published in Cell in 2014 also contains a compartment format which is shown below:

| | | | | | | | | |
|---|---|---|---|---|---|---|---|---|
| chr19 | 0 | 200000 | NA | 0 | . | 0 | 200000 | 255,255,255 |
| chr19 | 200000 | 500000 | B1 | -1 | . | 200000 | 500000 | 220,20,60 |
| chr19 | 500000 | 3800000 | A1 | 2 | . | 500000 | 3800000 | 34,139,34 |
| chr19 | 3800000 | 3900000 | B1 | -1 | . | 3800000 | 3900000 | 220,20,60 |
| chr19 | 3900000 | 5000000 | A1 | 2 | . | 3900000 | 5000000 | 34,139,34 |
| chr19 | 5000000 | 5600000 | B1 | -1 | . | 5000000 | 5600000 | 220,20,60 |

**Example annotation painting**

Choose the format of your annotation data for painting from the dropdown menu:

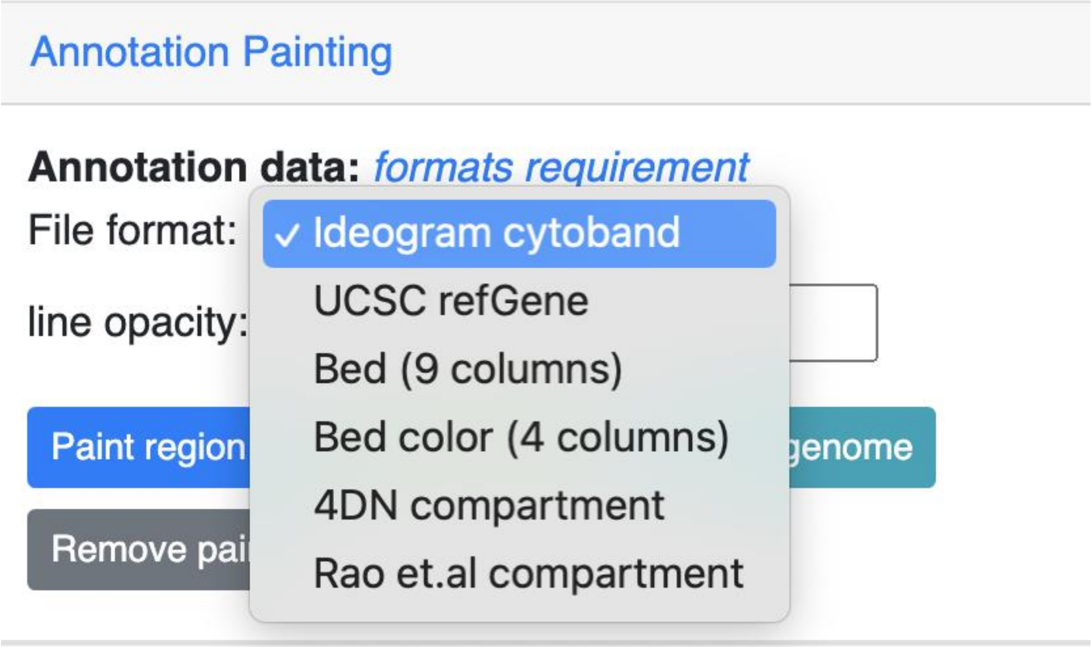

Then click a the paint button. If the format is not cytoband, an upload file button will appear.

**cytoband painting**

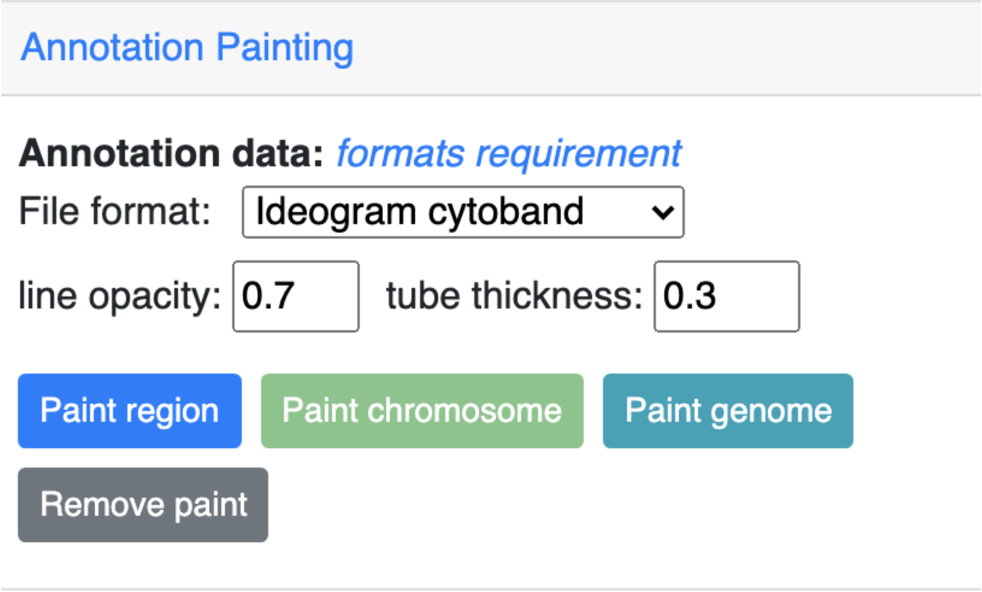

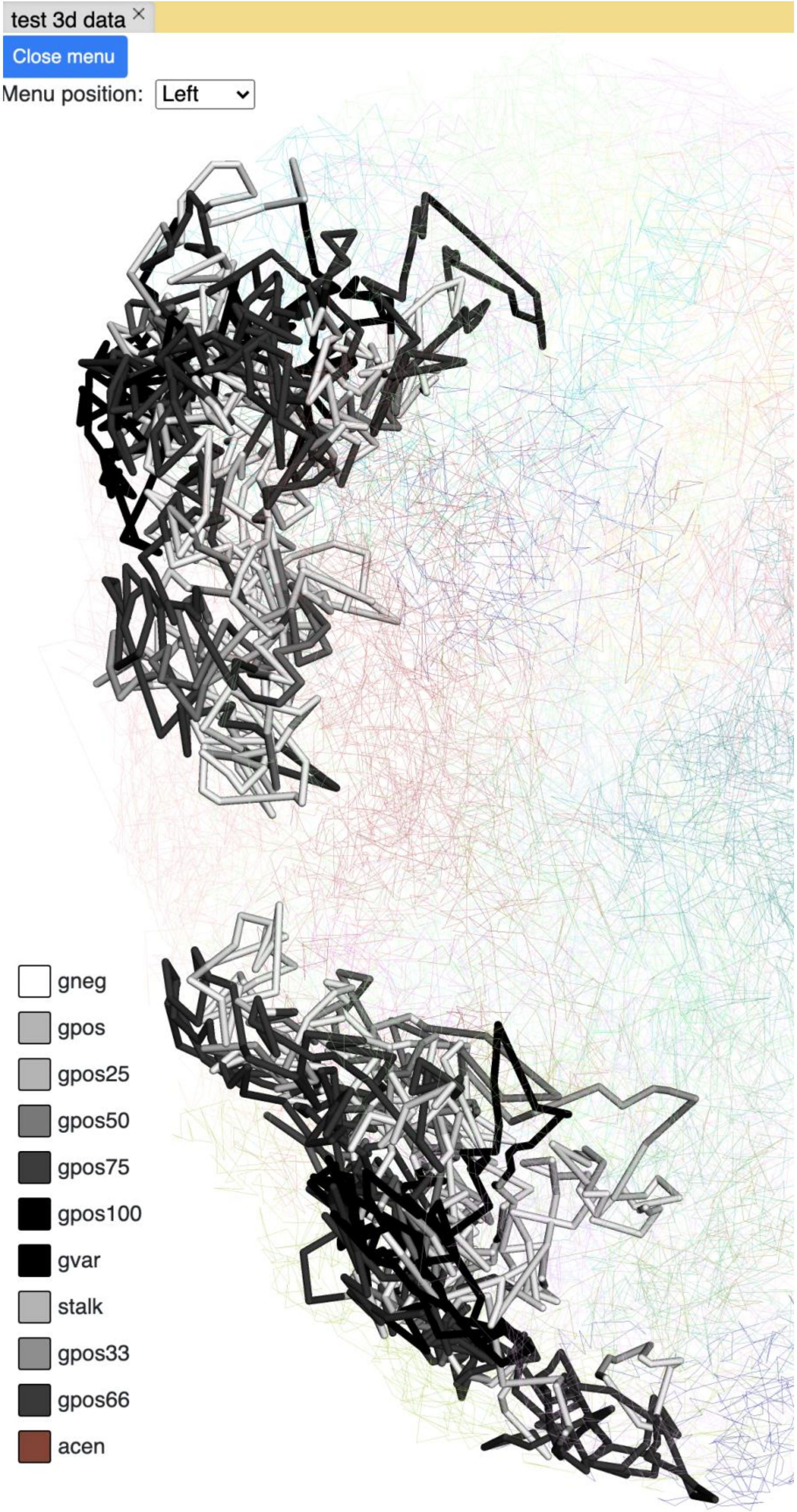
test 3d data
Close menu
Menu position:
Left
gneg
gpos
gpos25
gpos50
gpos75
gpos100
gvar
stalk
gpos33
gpos66
acen

**4DN compartment painting**

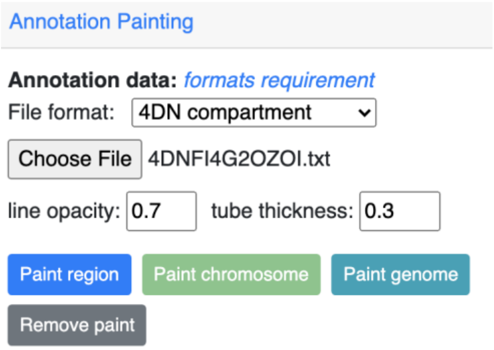

The screenshot below is an example using the compartment calls table mentioned above to paint a whole chromosome. The green part indicates compartment A and the red indicates compartment B. These colors can also be customized. The options are same as those for numerical painting.

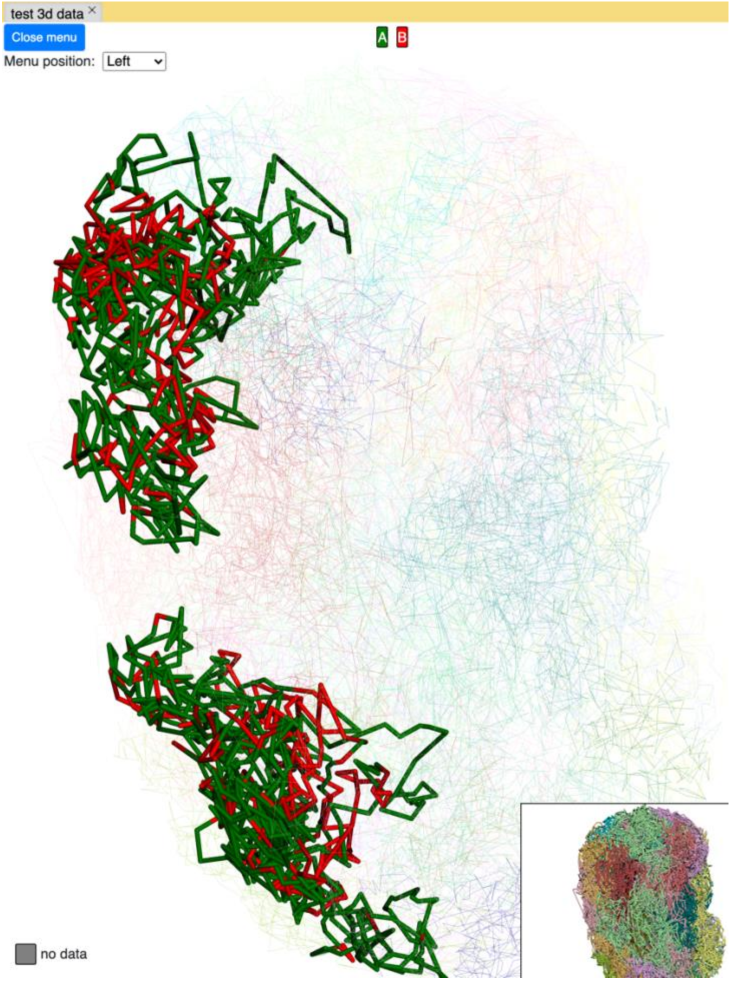

**chromHMM painting**

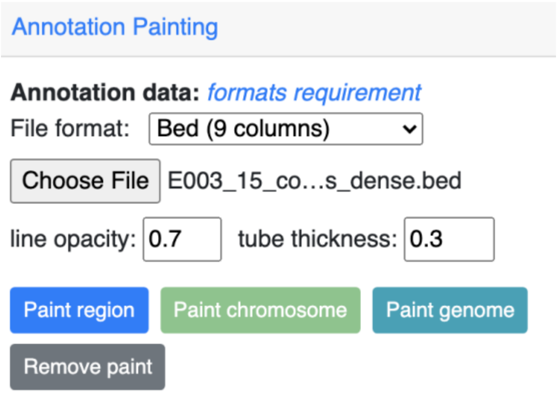

The screenshot below is an example using the chromHMM data from Roadmap project to paint a whole chromosome.

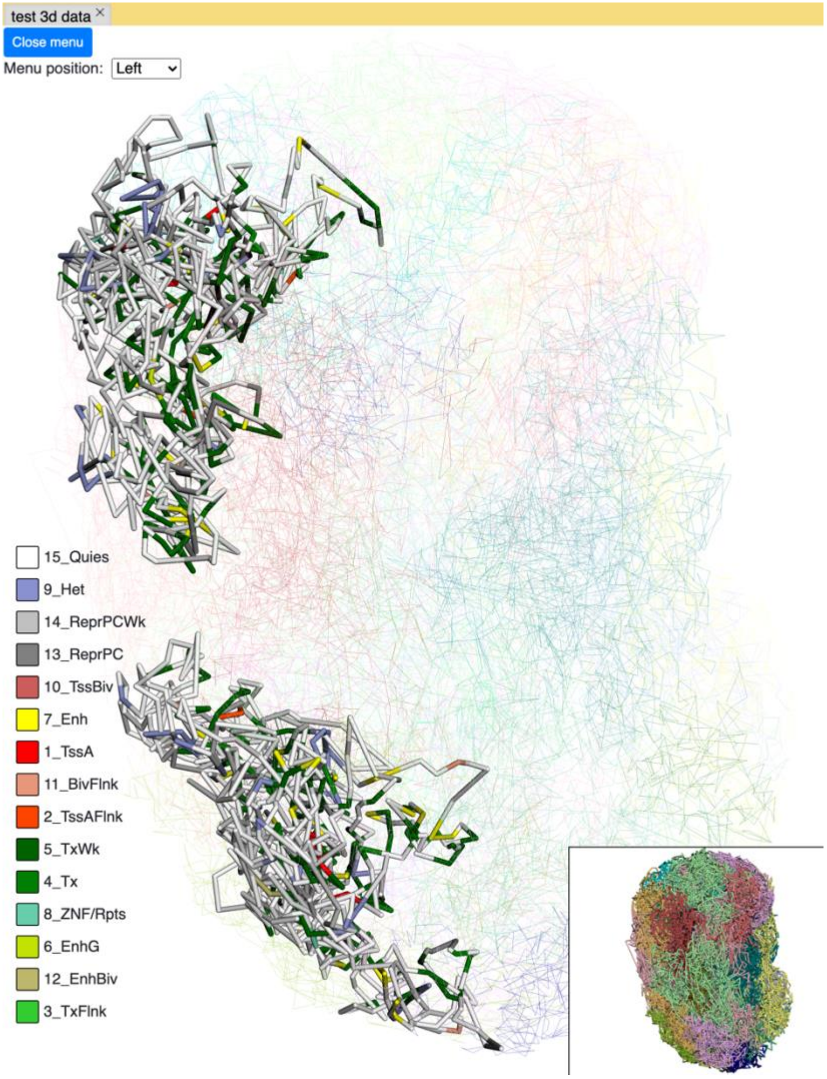

**Animations on 3D**

g3d format is designed to be a container file format. It might contain multiple models from haplotypes or different cells/samples; each model may also contain data at different resolutions. This example file contains 3D structure data from 3 different cells at different resolutions. When there are multiple models available, the 3D viewer can play an animation by looping over each model and displaying it as a frame. To view this example animation, add this example file as g3d track to the 3D browser:

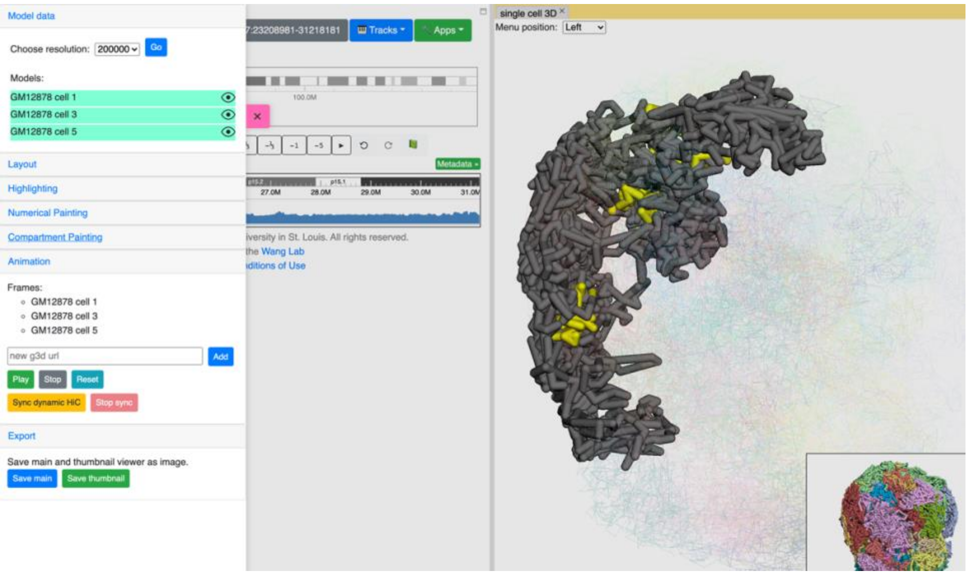

In the *Animation* section, click *Play* to start the animation and *Stop* to end it. The *Reset* button will reset the viewer to default view style.

**Export 3D images**

The 3D viewer can export the current view as image in png format for download. Simply click the save buttons under the *Export* section to download an image of the main and thumbnail viewers.

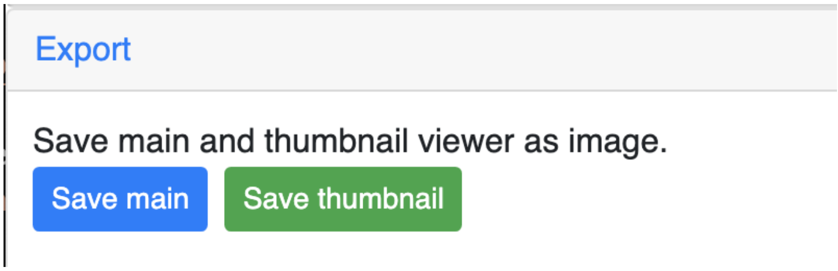
